# p16^High^ immune cell - controlled disease tolerance as a broad defense and healthspan extending strategy

**DOI:** 10.1101/2024.07.15.603540

**Authors:** Francisco Triana-Martinez, Alessandra Pierantoni, Daisy Graca, Veronica Bergo, Alexander Emelyanov, Bogdan B Grigorash, Shunya Tsuji, Sosuke Nakano, Laurent Grosse, Vesna Brglez, Pierre Marty, Jean Dellamonica, Albert J Fornace, Eirini Trompouki, Eiji Hara, Barbara Seitz-Polski, Dmitry V Bulavin

**Affiliations:** Institute for Research on Cancer and Aging of Nice (IRCAN); Université Côte d’Azur, INSERM; CNRS, Nice, France; Laboratoire d’Immunologie, Centre Hospitalier Universitaire de Nice, Nice, France; UR2CA - Unité de Recherche Clinique Côte d’Azur, Université Côte d’Azur (UCA), Nice, France; Department of Cellular and Molecular Immunology, Max Planck Institute of Immunobiology and Epigenetics, Freiburg, Germany; Faculty of Biology, University of Freiburg, Freiburg, Germany; International Max Planck Research School for Molecular and Cellular Biology (IMPRS-MCB), Freiburg, Germany; Research Institute for Microbial Diseases, Osaka University, Suita, Osaka, 565-0871, Japan; CHU Hôpital de l’Archet 1, Nice France; Service de Médecine Intensive Réanimation, CHU, Nice, France; Department of Oncology, Lombardi Comprehensive Cancer Center, Washington, DC; Department of Biochemistry and Molecular & Cellular Biology, Georgetown University Medical Center, Washington, DC

## Abstract

The ability of an organism to overcome infectious diseases has traditionally been linked to killing invading pathogens. Accumulating evidence, however, indicates that, apart from restricting pathogen loads, organismal survival is coupled to an additional yet poorly understood mechanism called disease tolerance. Here we report that p16^High^ immune cells play a key role in establishing disease tolerance. We found that the FDA-approved BNT162b2 mRNA COVID-19 vaccine is a potent and rapid inducer of p16^High^ immune subsets both in mice and humans. In turn, p16^High^ immune cells were indispensable for counteracting different lethal conditions, including LPS-induced sepsis, acute SARS-CoV-2 infection and ionizing irradiation. Mechanistically, we propose that activation of TLR7 or a low physiological activity of STING is sufficient to induce p16^High^ immune subset that, in turn, establishes a low adenosine environment and disease tolerance. Furthermore, containing these signals within a beneficial range by deleting MDA5 that appeared sufficient to maintain a low activity of STING, induces p16^High^ immune cells and delays organ deterioration upon aging with improved healthspan. Our data highlight the beneficial role of p16^High^ immune subsets in establishing a low adenosine environment and disease tolerance.

## Introduction

The general decline of physiological fitness and loss of tissue homeostasis with age is commonly associated with the onset and progression of multiple diseases, including cancer, diabetes, Alzheimer’s, osteoarthritis and many others^1^. Different mechanisms have been proposed to contribute to this natural phenomenon such as telomere attrition, mitochondrial dysfunction, and enhanced inflammation^2^. Importantly, many of these mechanisms can be directly linked to the induction of cellular senescence^3,4^, a well-characterized response that hinders cell cycle progression through the activation of the tumor suppressor genes *CDKN1A* (p21) or *CDKN2A* (p16)^5–7^. Substantial experimental evidence suggests that the accumulation of senescent cells is an important factor in age-related tissue deterioration^4,8^ as it is associated with the production of different molecules capable of restructuring the extracellular matrix, modifying the behavior of neighboring cells and systemically affecting the activity of immune system^9–11^. Despite these deleterious functions of senescent cells in the aging process, accumulating evidence supports cellular heterogeneity among p16^High^ cells with some mediating important homeostatic functions that have been identified during embryonic development as well as in adult skin, liver and lung^12–17^. This suggests that depending on the context, p16^High^ senescent cells could be either beneficial or detrimental. What defines either group remains however largely unknown.

The development of different genetic mouse models is now facilitating the further identification and characterization of p16^High^ cells *in vivo*^8,12,15,18,19^. Among the different p16^High^ subtypes, cells of the immune system, including T cells and macrophages, have been identified and further analysis revealed that some express additional markers of senescence such as enhanced senescence-associated β-galactosidase (SA-β-gal) activity and DNA damage^19–21^. Furthermore, the frequency of such cells increases significantly in animals during natural and accelerated aging, which may highlights their potential importance^21–23^. On the other hand, a modest or even transient activation of p16, as well as excessive lysosomal activity (and thus higher SA-β-gal activity) in phagocytic cells such as macrophages has been observed under different conditions^19,20^. Whether such activation indeed reflects classical pathways of senescence activation is unclear.

In our current study, we used a genetic mouse model to trace cells with high expression of p16 *in vivo*. We found that the p16^High^ program was activated during aging not only in long-lived macrophages and T cells, but in all the immune subsets analyzed. Our detailed analysis of T cells and tissue-resident macrophages as well as the use of a genetic model for selective ablation of p16^High^ cells, allowed us to determine that p16^High^ immune cells play an important regulatory functions *in vivo*. These functions were further critical for animal survival after severe inflammation and tissue damage. While the ability of an organism to overcome infectious diseases has traditionally been linked to killing invading pathogens, evidence indicates that, apart from restricting pathogen loads, organismal survival is coupled to an additional yet poorly understood mechanism called disease tolerance^24,25^. Here we argue that induction of p16^High^ immune cells is a key mechanism in establishing disease tolerance.

## Results

### p16^High^ immune cells accumulate in different tissues with age

Using the transgenic murine model p16-Cre/R26-mTmG, we selectively identified cells with high expression of p16 (henceforth designated p16^High^ cells) by detecting the expression of an EGFP reporter gene^12^. We performed flow cytometry-based analysis of p16^High^ cells resident in different murine tissues at 2, 12, and 18 months-of-age, including the bone marrow, peripheral blood, spleen, peritoneal cavity, liver and the stromal-vascular fraction (SVF) of abdominal fat. Next, we determined the percentage of cells expressing high levels of p16 within populations of cells containing markers that are enriched in different immune cell subtypes, including CD3 (T cells), B220 (B Cells), CD11c (dendritic cells), Ly6C (monocytes), Ly6G (neutrophils), and F4/80 (macrophages). Overall, we observed a relative increase in the numbers of p16^High^ cells with age in all the tested immune subsets (Figure 1A). Further analysis of different immune subsets revealed their significant contribution to the total number of p16^High^ cells in different tissues ranging from 35%–40% for SVF in abdominal fat and liver, to more than 80% in the spleen (Figure 1B). Thus, p16^High^ cells are present both in very short-lived (Ly6G^+^, neutrophils) and very long-lived (F4/80^+^, tissue-resident macrophages) immune subsets and constitute a significant fraction of the total p16^High^ population, thus potentially representing senescent cells in different tissues of aging animals.

**Figure 1.**
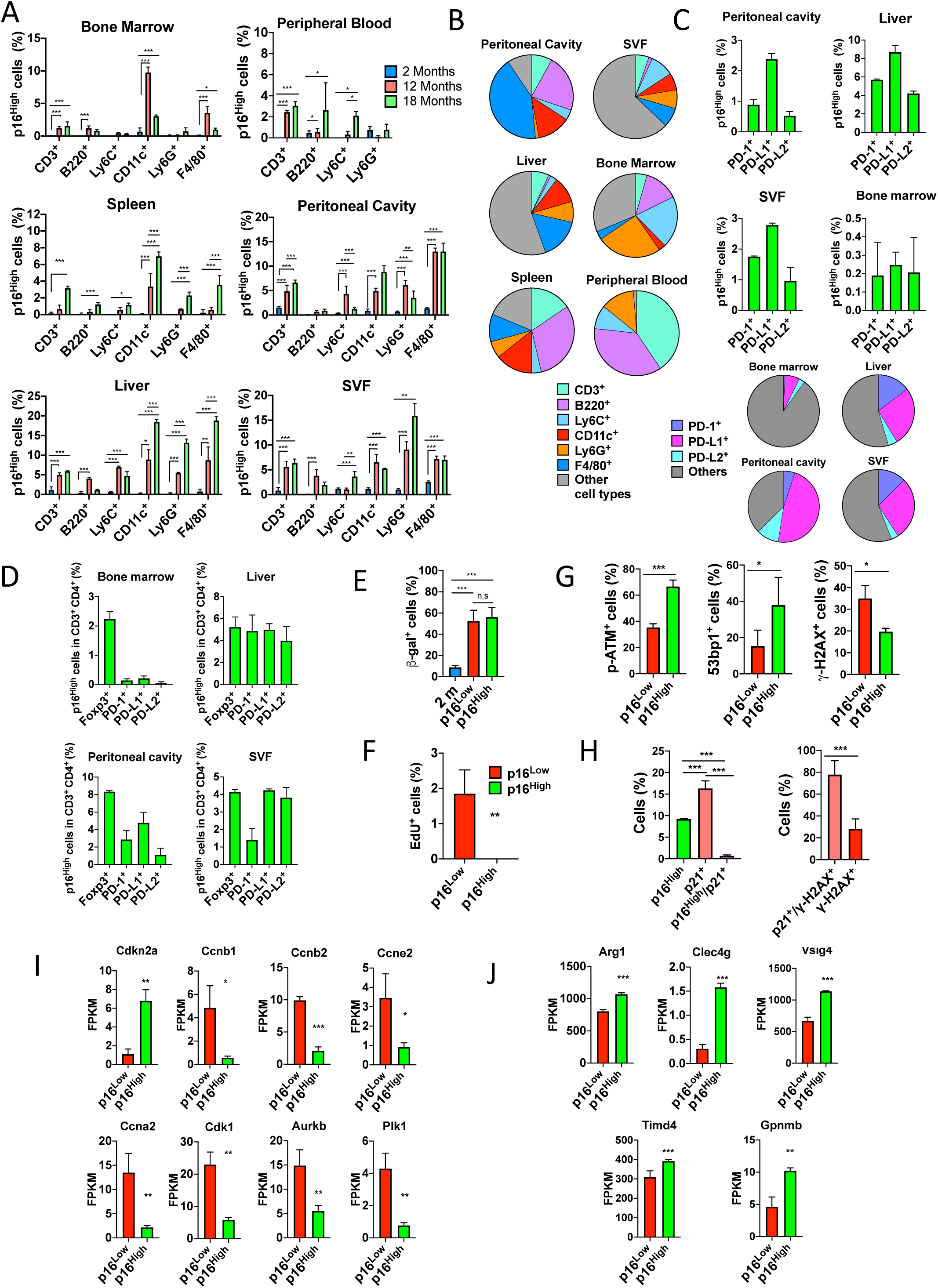
Characterization of p16^High^ immune cells in different tissues. **A.** Analysis of the fraction of p16^High^ cells in different populations of the immune cells. Single cell suspensions were prepared from peritoneal cavity, spleen, liver, stromal-vascular fraction of abdominal fat (SVF), bone marrow, and peripheral blood from 2-, 12- and 18-month-old p16-Cre/R26-mTmG mice. Cell suspensions were stained with fluorescent-conjugated antibody against the corresponding markers of immune cells (CD3 for T lymphocytes, B220 for B lymphocyte, Ly6C for monocytes, Ly6G for neutrophils, CD11c for dendritic cells and F4/80 for tissue-resident macrophages) and analyzed by flow cytometry. Independent repeats *n = 3*. A representative experiment is shown. Data are mean ± S.D. Statistical significance was analyzed using ANOVA plus Tukey post hoc test. *p<0,05; **p<0,01 and ***p<0,001. **B.** Abundance and distribution of p16^High^ cells from the previously analyzed tissues in 18-month-old mice. Each pie chart represents the total number of p16^High^ cells and how they are distributed in the different cellular compartments. **C.** Percentage of p16^High^ cells in the populations expressing PD-1, PD-L1 and PD-L2. Peritoneal, liver, bone marrow and abdominal fat SVF single cell suspensions from 16-month-old mice were stained using fluorescent-conjugated antibody against PD-1 and PD-L1 and analyzed by flow cytometry. Bar graph represents the percentage of p16^High^ in populations expressing PD-1 and PD-L1. Pay charts show the distribution of the total p16^High^ cells expressing PD-1, PD-L1 or PD-L2. Independent repeats *n = 3*. Data are mean. **D.** p16^High^ T lymphocytes express PD-1, PD-L1, PD-L2 and Foxp3. The percentage of p16^High^ cells was determined in T lymphocytes obtained from peritoneal cavity, liver, bone marrow and abdominal fat SVF defined as CD3^+^ CD4^+^ PD1^+^, CD3^+^ CD4^+^ PD-L1^+^, and CD3^+^ CD4^+^ Foxp3^+^ in 16-month-old mice. Data are mean ± S.D. **E.** Senescence-associated β-galactosidase activity was analyzed in 2-month-old and 12-month-old p16^High^ and p16^low^ F4/80-positive macrophages. Pictures were made with bright field – fluorescence microscope. The percentage of positive cells was calculated. Data are mean ± SD. Significant differences were determined by t-test. Independent repeats *n = 6*. *p<0,05; **p<0,01 and ***p<0,001. **F.** p16^High^ peritoneal macrophages are not proliferative. Peritoneal macrophages from 12-month-old p16-Cre/R26-mTmG mice were incubated with 5-ethynyl-2’-deoxyuridine (EdU) for 24 h. The percentage of EdU positive cells was analyzed in p16^High^ and p16^Low^ populations. Data are mean ± SD. Significance was analyzed by t-test. **p<0,01. **G.** Immunostaining of different markers of senescence. Peritoneal macrophages from 12-16 month-old p16-Cre/R26-mTmG mice were stained for senescence markers ATM, 53bp1, and ϒ-H2AX. Pictures were obtained by high-definition or confocal microscopy. Cells were counted and the percentages of positive cells for p16^High^ and p16^low^ populations were calculated. Data are mean ± SD. Significant differences were determined by t-test. *p<0,05; **p<0,01 and ***p<0,001. **H.** Peritoneal macrophages from 12-16 month-old p16-Cre/R26-mTmG mice were stained for p21 and ϒ-H2AX. Pictures were obtained by high-definition or confocal microscopy. Cells were counted and the percentage of p16^High^, p21^+^ or double positive cells were determined (left panel). The percentage of positive cells for ϒ-H2AX^+^, and ϒ-H2AX^+^ in p21^+^ (right panel) were determined. Independent repeats *n = 6.* Data are mean ± SD. Significant differences were determined by t-test. ***p<0,001. **I.** Peritoneal F4/80^+^ macrophages from 12-month-old p16-Cre/R26-mTmG mice were isolated using magnetic cell sorting. Then, p16^High^ and p16^low^ populations were separated by fluorescence-activated cell sorting (FACS). RNA expression of both populations was analyzed by RNA sequencing. Kegg pathway analysis of downregulated genes from RNA-seq data set reveals a set of clusters of genes related with the negative control of cell cycle. Individually analyzed genes are shown. Fragments Per Kilobase of transcript sequence per Millions base pairs sequenced (FPKM) were analyzed by t-test. *n = 3* biologically independent samples. Data are mean ± SD. *p<0,05; **p<0,01 and ***p<0,001. Representative 8 genes are shown. **j.** Kegg pathway analysis of upregulated genes from RNA-seq data set reveals a set of clusters related with the negative control of immune cells and T-cells. Immunosuppressive and regulatory representative genes were individually analyzed and shown. Data are mean ± SD. Differences were established with t-test. **p<0,01 and ***p<0,001.

Recent data showed an enrichment of PD-1–PD-L1 markers in senescent cells^26^; therefore, we next determined the fraction of p16^High^ cells among the total PD1^+^, PD-L1^+^ and PD-L2^+^ cells in bone marrow, liver, peritoneal cavity, and SVF in 16-month-old mice. We found that the contribution of p16^High^ cells in these populations varied significantly depending on the tissue analyzed, from 0.2 % in the bone marrow to 6-8 % in the liver (Figure 1C). Analysis of the fraction of PD1^+^, PD-L1^+^ and PD-L2^+^ cells in the total number of p16^High^ cells revealed a wide range from a small (bone marrow) to a more significant (peritoneum) contribution (Figure 1C, pie charts). Thus, the contribution of p16^High^ cells to the total number of PD1^+^, PD-L1^+^ and PD-L2^+^ cells and vice versa varies significantly among different tissues, with PD-L1^+^ cells representing a relatively higher proportion of the p16^High^ cell population (Figure 1C, bottom charts).

As PD-1, PD-L1 and PD-L2 are frequently found on immunosuppressive T cells, we next determined the expression of p16 in these subtypes and also extended our analysis to regulatory T cells (Tregs) as another suppressive immune subset^21,27^. For this purpose, we analyzed the fractions of CD3^+^ CD4^+^ PD-1^+^, CD3^+^ CD4^+^ PD-L1^+^, CD3^+^ CD4^+^ PD-L2^+^and CD3+ CD4^+^ FoxP3^+^ cells in the p16^High^ cell populations (Figure 1D). We found strong enrichment of T cells expressing PD-1, PD-L1 but also regulatory FoxP3 markers in the p16^High^ cell populations, suggesting that a significant fraction of the total PD1^+^ and PD-L1^+^ cells expressing high levels of p16 in any given tissue (Figure 1C) are indeed T cells. Further analysis of different subsets among the p16^High^ T cells revealed a strong enrichment for Tregs. By contrast, PD-L2^+^ cells were less enriched inside the p16^High^ T cells (Figure 1D).

Overall, our analysis showed that a noticeable fraction of p16^High^ T cells express markers of regulatory subsets, and thus potentially could play an important role in negative regulation of tissue inflammation and damage through well-established immunosuppressive and tissue remodeling activities of Tregs^27–29^.

### p16^High^ peritoneal macrophages express few markers of senescence and regulatory

Tissue-resident macrophages, in comparison to other immune subtypes, are of embryonic origin and maintained for the entire life span of the organism^30^. Based on their longevity, we speculated that tissue-resident macrophages could be the best immune cell candidates to express multiple markers of senescence in addition to the high levels of p16 as an indicator of full senescence. To investigate this hypothesis in more detail, we focused on F4/80^+^ peritoneal tissue-resident macrophages^31^. Previous studies have shown that macrophages exhibit a modest increase in expression levels of p16 and SA-β-gal, both putative markers of senescence, with age and in response to the presence of senescent cells in the peritoneal cavity^19,20^. To investigate peritoneal F4/80^+^ macrophages further, we analyzed the SA-β-gal activity in these cells isolated from 2- and 12-month-old p16-Cre/R26-mTmG mice. We found that while SA-β-gal activity was indeed significantly higher in the cells isolated from 12-month-old mice when compared to those isolated from to 2-month-old mice, surprisingly, we found no difference in the levels of activity between the p16^High^ (EGFP expressing cells) and p16^Low^ (tdTomato expressing cells) populations (Figure 1E). Importantly, our analysis of proliferation using BrdU labeling confirmed the absence of proliferation by p16^High^ peritoneal F4/80^+^ macrophages, which is a feature of a cell cycle arrest (Figure 1F). In addition, we found a reduction in the migratory capacity of p16^High^ F4/80^+^ macrophages, while there was no difference in phagocytic activity between the p16^High^ and p16^Low^ populations. Analysis of additional putative senescence markers revealed an increase in the percentage of both p-ATM^+^ and 53bp1^+^ cells, while γH2Ax was reduced in the p16^High^ population (Figure 1G). To explore further this unexpected reduction in γH2Ax+ cells in p16^High^ population, we evaluated the expression of p21 protein, another putative marker of senescence. Double-positive p21^+^/p16^+^ cells were infrequent and further analysis revealed that the majority (approximately 80%) of the p21^+^ cells were positive for γH2Ax (Figure 1H). Thus, our results showed the presence of two populations of peritoneal F4/80^+^ macrophages that carry senescent markers in aging mice - one, which is positive for p16 and p-ATM/53Bp1, and another, which is positive for p21 and γH2Ax - with both populations expressing SA-β-gal.

To gain a better understanding of the changes that accompany p16 activation, we performed RNA-sequencing (RNA-seq) on p16^High^ and p16^Low^ F4/80+ peritoneal macrophages from 12-month-old animals. First, we confirmed the upregulation of p16 mRNA in p16^High^ cells and subsequent KEGG pathway analysis revealed that the most represented clusters of downregulated genes were related to cell cycle arrest and DNA replication (Figure 1I). Next, we manually curated the list of genes related to the senescence-associated secretory phenotype (SASP). Surprisingly, the number of over-expressed factors associated with the SASP was limited in p16^High^ cells, while none of the classical inflammatory factors were over-expressed. KEGG analysis of the upregulated genes revealed a set of genes related to the regulatory functions of immune cells including the *Arg1*, *Clec4g*, *Vsig4*, *Timd4* and *Gpnmb* genes (Figure 1J). Thus, our analysis of p16^High^ peritoneal F4/80^+^ macrophages revealed that these cells express high levels of p16, resulting in profound cell cycle arrest, as well as the expression of regulatory factors that exhibit both immunosuppressive and tissue remodeling activities such as *Arg1 and Clec4g* ^32–34^. Overall, our macrophage analysis together with the observed expression of several regulatory and immunosuppressive markers by p16^High^ T cells (Figure 1C&D) suggested that p16 activation is accompanied by the appearance of different regulatory and immunosuppressive immune subsets.

### p16^High^ immune cells arise in young animals in response to inflammation and tissue damage

Since our data in T cells (Figure 1C&D) and macrophages (Figure 1J) pointed towards regulatory nature of p16^High^ immune cells, next we seek further confirmation of these observations at the tissue level. We speculated that removal of p16^High^ immune cells with subsequent analysis of tissue pro- and anti-inflammatory profiles would allow us to define more precisely their role. This is something that would be difficult to achieve in an old organism due to present of different p16^High^, including numerous types of senescent cells in multiple tissues. Thus, we seek the conditions for the accumulation of p16^High^ immune cells in young organisms so their clearance would provide a clear confirmation of their functions at a tissue level. Aging is accompanied by a noticeable intestinal deterioration with the development of a so-called “leaky gut”^35^. If the process spreads further, this increased intestinal wall permeability can trigger a defensive inflammatory response in both the peritoneum and the liver^12,36^. To mimic an age-induced deterioration of the intestine, we employed a model of dextran sodium sulfate (DSS)-induced colitis^37^ in 2-month-old p16-Cre/R26-mTmG mice and analyzed the accumulation of p16^High^ immune subsets in the peritoneal cavity and liver by flow cytometry. We found an increase in the percentage of p16^High^ cells in different immune subsets both in the peritoneum and liver (Figure 2A). Analysis of absolute numbers of p16^High^ cells revealed that their largest fraction consisted of CD3^+^ T cells and F4/80^+^ macrophages (Figure 2B). Among the CD3^+^ T cells, we found a significant enrichment of p16^High^ cells expressing the regulatory marker Foxp3, as well as PD1 and PD-L1 (Figure 2C&D). Thus, DSS-induced colitis was confirmed to trigger an efficient accumulation of p16^High^ immune subsets in young mice. We next investigated the impact of their genetic ablation using a diphtheria toxin A (DTA)-dependent ablation mouse line, in which p16^High^ cells are selectively eliminated through activation of the Cre recombinase (p16-Cre/R26-DTA). p16-Cre/R26-mTmG and p16-Cre/R26-DTA mice were either mock- or DSS-treated and RNA was isolated from peritoneal cells and liver for qPCR analysis. First, we confirmed an increase in the level of p16 mRNA in both tissues after DSS treatment of p16-Cre/R26-mTmG mice, something that was fully abrogated in p16-Cre/R26-DTA mice. The *Arg1* and *Il10* genes have been implicated in both immunosuppression and tissue remodeling^38,39^ and their analysis showed strong accumulation in both peritoneal and liver cells after DSS treatment of p16-Cre/R26-mTmG, but not in p16-Cre/R26-DTA mice (Figure 2E&F). In contrast, we observed marked accumulation of the pro-inflammatory genes *IL1b* and *IL6* in the peritoneal cavity after selective ablation of p16^High^ immune cells (Figure 2E). As such, our analysis directly confirmed that at the tissue level, p16^High^ immune cells have an important regulatory functions and their depletion under certain conditions can trigger an enhanced inflammatory and tissue damage responses.

**Figure 2.**
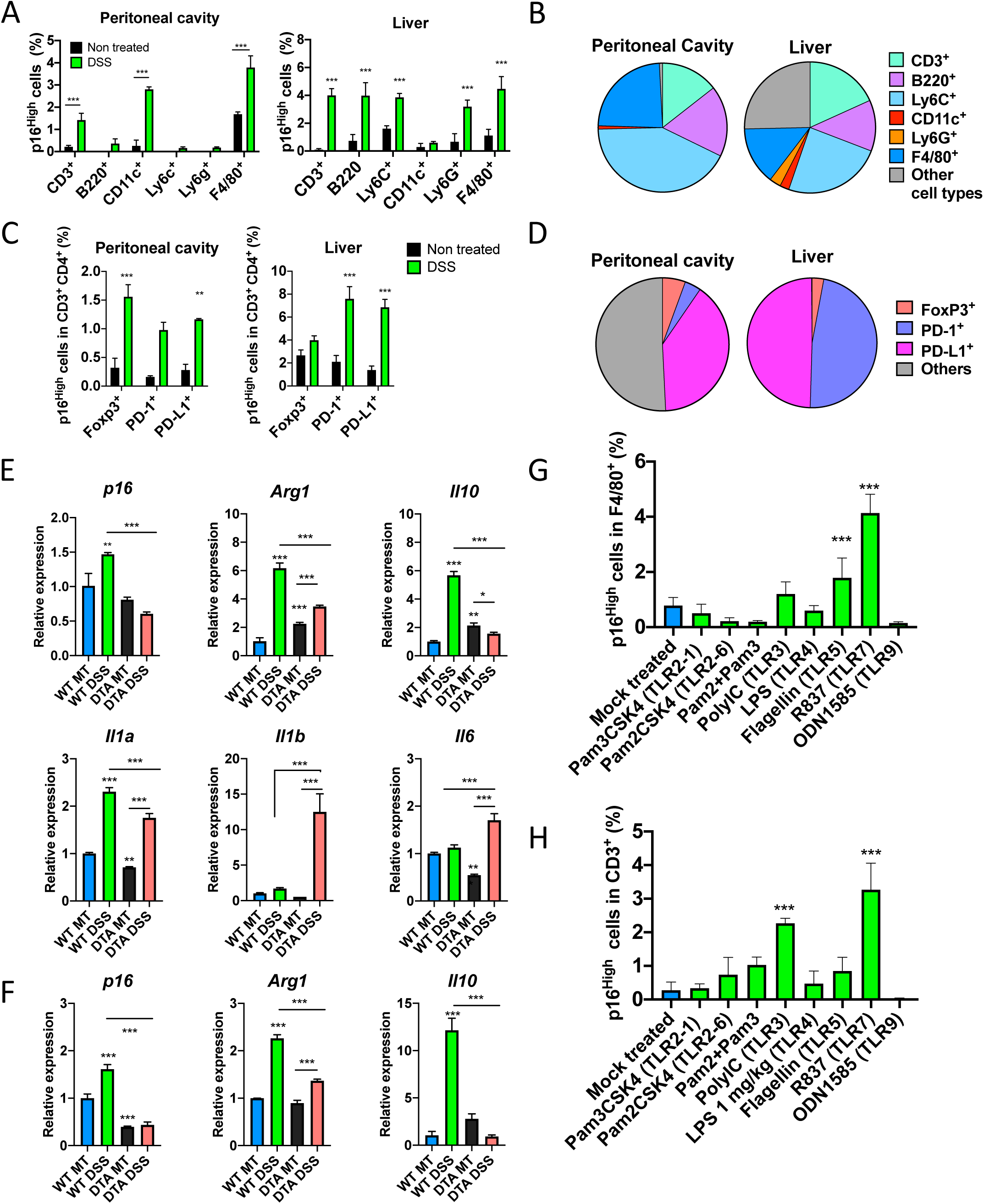
p16^High^ immune cells arise in young animals in response to inflammation and tissue damage. **A-B**. Dextran sodium sulphate (DSS) induces p16^High^ cells *in vivo* in young mice. 2-3 month-old p16-Cre/R26-mTmG mice were treated with DSS 2.5% dissolved in drinking water for 7 days and the percentage of p16^High^ cells in different populations of immune system was determined. Single cell suspensions from peritoneal cavity and liver were stained with fluorescent conjugated antibody and analyzed by flow cytometry. Abundance and distribution of p16^High^ cells in the previously analyzed tissues (panel A) were determined and are shown as pie charts (**B**). *n = 4* biologically independent samples. Data are mean ± SD. Statistical significance was determined using ANOVA plus Dunnett post hoc test. **p<0,01 and ***p<0,001. **C-D**. p16^High^ T lymphocytes express higher level of Foxp3, PD-1, and PD-L1 after DSS treatment. The percentage of p16^High^ cells was determined in T lymphocytes obtained from peritoneal cavity and liver defined as CD3^+^ CD4^+^ PD1^+^, CD3^+^ CD4^+^ PD-L1^+^, and CD3^+^ CD4^+^ Foxp3^+^ I in 12-month-old mice. Pie charts show abundance and distribution of Foxp3, PD-1, and PD-L1 in the total CD3^+^ p16^Hgh^ population (**D**). Data are mean. **E-F**. Different pattern of gene expression in the absence of p16^High^ cells. Control and p16-Cre/R26-DTA (DTA) were treated with 2.5% DSS for 7 days and RNA from peritoneal cells (**E**) and liver (**F**) were isolated. RNA was analyzed by quantitative SYBR-green based PCR (qPCR) to determine the level of expression of different genes. Data are mean +/- S.D. Independent repeats *n = 3.* Difference between groups was analyzed using ANOVA test and Tukey post hoc test. *p<0,05; **p<0,01 and ***p<0,001. **G-H**. Specific TLRs induce p16^High^ state *in vivo*. 2-3 months p16-Cre/R26-mTmG mice were treated intraperitoneal (I.P) with a panel of agonist to different toll-like receptors (TLRs) (see Materials and Methods section for detailed treatments). 48 h later, peritoneal cells were stained with fluorescent-conjugated antibodies against F4/80 (**G**) and CD3 (**H**). The percentage of p16^High^ cells was determined by flow cytometry. Data are mean ± SD. *n = 3* biologically independent samples. Statistical significance were determined using ANOVA plus Dunnett post hoc test. *p<0,05 and ***p<0,001.

To gain potential mechanistic insights, we explored the ability of Toll-like receptor (TLR) stimulation, which also occurs in a DSS model of colitis^37,40^, to induce p16^High^ immune subsets. TLRs are the first-line sensors and responders to possible harmful factors including different pathogens, in the surrounding environment. These receptors act as defense and homeostatic regulators by recognizing both pathogen-associated molecular patterns (PAMPs) and damage-associated molecular patterns (DAMPs)^41^. To systematically address the role of different TLRs in induction of p16^High^ immune cells, we administered different TLR agonists to 2-month-old p16-Cre/R26-mTmG mice and analyzed p16^High^ immune cells in the peritoneal cavity. We found that agonists of TLR7 (R837) and to a lesser extent TLR5 (flagellin) were the best inducers of p16^High^ macrophages (Figure 2G), while agonists of TLR1-2 (Pam3CSK4) and TLR7 (R837) were the best inducers of p16^High^ T cells (Figure 2H). Overall, these data suggested that induction of p16^High^ immune cells could be a part of a physiological response to tissue damage and inflammation and can be triggered by different TLRs receptors. Among the TLRs, TLR7 appeared to be strongest inducer of p16^High^ macrophages and T cells, both of which are long-lived immune subsets.

### The BNT162b2 mRNA COVID-19 vaccine induces regulatory p16^High^ immune cells

Since TLR7 receptor activation efficiently induced different p16^High^ immune subsets, we focused on this signaling pathway for further analysis. TLR7 is located intracellular, primarily in endosomes and detects single-stranded RNA targets, including those from different viruses such as HIV, HCV, and SARS-CoV-2 ^42,43^. For further analysis, we focused on the BNT162b2 mRNA COVID-19 vaccine as an only FDA-approved TLR7 agonist, which has also been reported to activate TLR8 and TLR9 expressed on antigen-presenting cells. We assessed the efficiency with which this vaccine induced different p16^High^ immune subsets. To do so, we intraperitoneal injected 2-month-old p16-Cre/R26-mTmG mice with BNT162b2 and investigated the induction of p16 in different immune cells at days 2 and 15. We found that BNT162b2 efficiently induced p16^High^ immune cell subtypes both in the peritoneum (Figure 3A&B) and liver, but also in CD31-positive liver sinusoid endothelial cells (LSECs), previously described as an abundant p16^High^ cell subset during aging^12^, which was observed as early as day 2 and lasted beyond day 15 after treatment. Further analysis of p16 induction in T cell subsets revealed an increase in cell populations positive for regulatory Foxp3, PD-1 and PD-L1 in the peritoneum (Figure 3C) and for Foxp3 in the liver after vaccine treatment.

**Figure 3.**
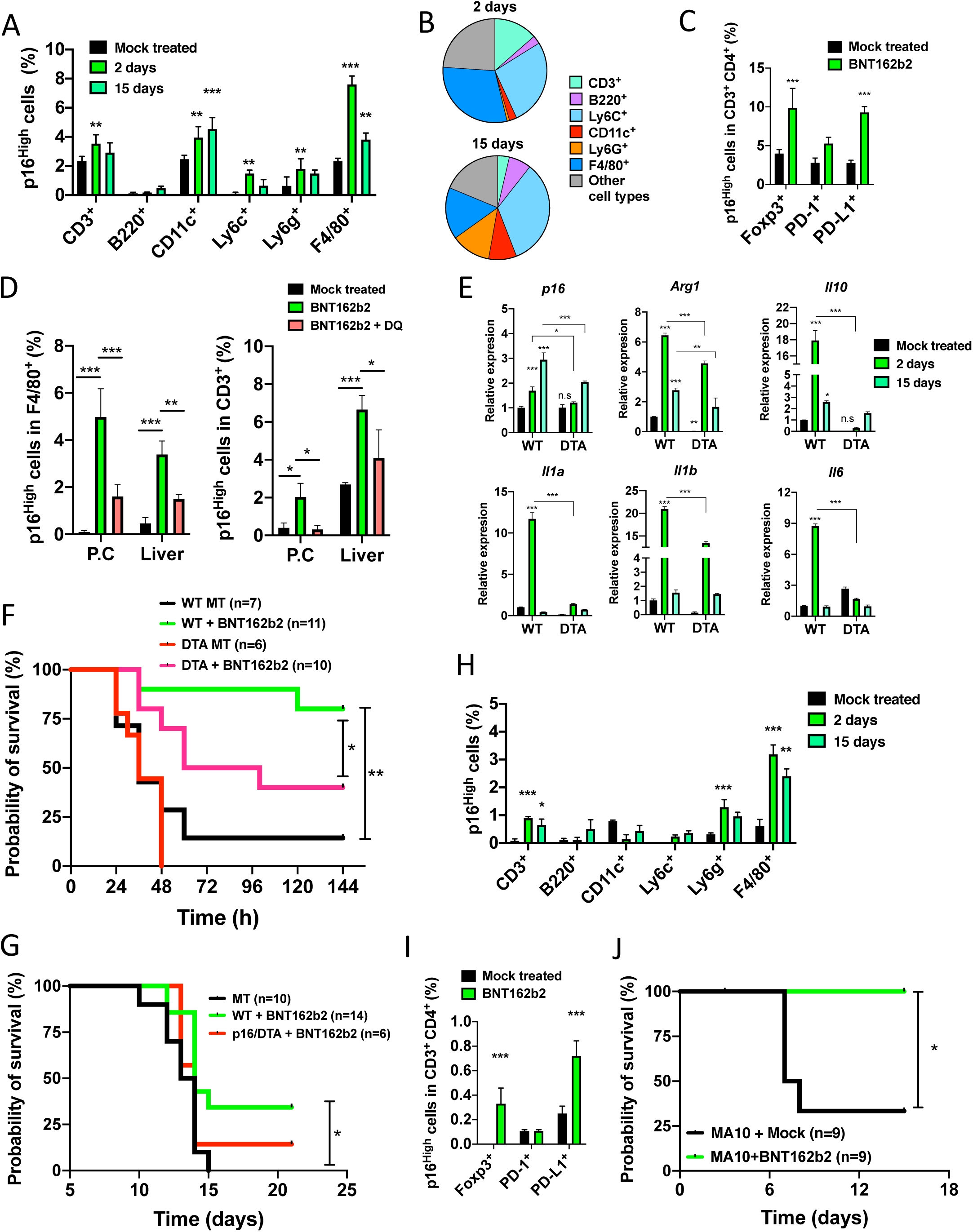
The BNT162b2 mRNA COVID-19 vaccine induces p16^High^ immune cells and disease tolerance to protect against severe inflammation and tissue damage. **A-B**. BNT162b2 mRNA COVID-19 vaccine induces p16^High^ cells *in vivo*. 2-3 month-old p16-Cre/R26-mTmG mice were treated intraperitoneal (I.P) with 5 µg of BNT162b2 mRNA COVID-19 vaccine. After 2 and 15 days the percentage of p16^High^ cells was determined in different immune subsets in peritoneal cavity (**A**). Abundance and distribution of different p16^High^ cells in analyzed tissues. Each pie chart represents the total number of p16^High^ cells and how they are distributed in different cellular compartments (**B**). Data are mean ± SD. *n = 4* biologically independent samples. Statistical significance was determined using ANOVA plus Dunnett post hoc test. **p<0,01 and ***p<0,001 **C.** The percentage of p16^High^ positive cells in different populations of T cells was determined at the same time than experiment shown in A. Data are mean ± SD. Statistical significance were determined using ANOVA plus Dunnett’s post hoc test ***p<0,001. **D.** BNT162b2 vaccine induces p16^High^ cells that are sensitive to senolytic treatment. 2-3 month-old p16-Cre/R26-mTmG mice were treated I.P with 5 µg of BNT162b2 vaccine. 5 days later, mice were either treated orally with a senolytic combination of dasatinib (5 mg/kg) and quercetin (50 mg/kg) or vehicle (Mock). After 24h mice were euthanized and single cell suspensions from peritoneal cavity and liver were stained with conjugated antibodies against F4/80 and CD3. Cells suspensions were analyzed by flow cytometry and the percentage of p16^High^ cells in each population was determined. *n = 3* biologically independent samples. Differences between groups were analyzed using ANOVA test and Tukey post hoc test. *p<0,05 and ***p<0,001. **E.** Genetic ablation of p16^High^ cells modifies the expression of inflammatory genes after BNT162b2 treatment. Wild type and p16-Cre/R26-DTA (DTA) were treated with 5 µg of BNT162b2 vaccine. After 2 and 5 days, RNA from peritoneal cells was isolated. RNA was analyzed by qPCR to determine the level of expression of inflammatory genes. Data are mean +/- S.D. Differences between groups were analyzed using ANOVA test and Tukey post hoc test. *p<0,05; **p<0,01 and ***p<0,001. **F.** p16^High^ cells protect against acute inflammation. Wild type and p16-Cre/R26-DTA (DTA) were either treated with 5 µg of BNT162b2 vaccine or saline (Mock). After 5 days, animals were injected with lipopolysaccharides (LPS) from Escherichia coli O55:B5 (LPS) 40 mg/kg. Animals were euthanized immediately once they reached the limit point of physical deterioration. Difference between groups was analyzed using Gehan-Breslow-Wilcoxon test. *p<0,05 and **p<0,01. *n* used for each condition is showed in the plot. **G.** p16^High^ cells protect against Ionizing Irradiation. Wild type and p16-Cre/R26-DTA (DTA) were treated with either 5 µg of BNT162b2 vaccine or saline (Mock). Animals were exposed to 8 Gy of γ-irradiation. Animals were observed daily and euthanized immediately once they reached the limit point of physical deterioration. Difference between groups was analyzed using Gehan-Breslow-Wilcoxon test. *p<0,05. *n* used for each condition is showed in the plot. **H.** BNT162b2 vaccine induces p16^High^ cells in the lung. 2-3 month-old p16-Cre/R26-mTmG mice were treated I.P with 5 µg of BNT162b2 vaccine. 2 and 15 days later, mice were euthanized and the percentage of p16^High^ immune cells was determined in the lung. Single cells suspensions from lung were stained with fluorescent conjugated-antibody against different markers of immune cells and analyzed by flow cytometry. Data are mean ± SD. Statistical significance were determined using ANOVA plus Dunnett post hoc test. **p<0,01 and ***p<0,001. *n = 3* biologically independent samples. **I.** BNT162b2 vaccine increases p16^High^ cells and induces Foxp3^+^, PD-1^+^ and PD-L1^+^ in CD3^+^ CD4^+^ populations. 2-3 month-old p16-Cre/R26-mTmG mice were treated I.P with 5 µg of BNT162b2 vaccine and after 5 days, single cell suspensions were stained with fluorescent-conjugated antibodies and the levels of CD3^+^ CD4^+^ PD1^+^, CD3^+^ CD4^+^ PD-L1^+^, and CD3^+^ CD4^+^ Foxp3^+^ populations and the percentage of p16^High^ cells were analyzed by flow cytometry. Data are mean ± SD. Statistical significance were determined using ANOVA plus Dunnett post hoc test. ***p<0,001. **J.** 2-3 months Balb/c mice were either treated with BNT162b2 vaccine or mock-treated and 3 days later were infected with a mouse-adopted MA10 virus. Analysis of probability of survival in experimental group as assessed, dead and mice that lost 70% and more body weight were considered as deceased. Difference between groups was analyzed using Gehan-Breslow-Wilcoxon test. *p<0,05. *n* used for each condition is showed in the plot.

Having previously found that the liver p16^High^ F4/80^+^ macrophages in 12-month-old mice are sensitive to a senolytic cocktail of dasatinib and quercetin (DQ)^12^, we next assessed the effects of BNT162b2 with and without subsequent DQ treatment (Figure 3D). We found that DQ efficiently reduced the numbers not only of p16^High^ macrophages, but also of T cells in both the peritoneum and liver in mice treated with BNT162b2. Thus, we confirmed that DQ treatment has a broad inhibitory activity on different p16^High^ immune subsets *in vivo*.

To understand the significance of BNT162b2-induced accumulation of p16^High^ immune cell subsets on the expression of inflammatory factors, next we treated 2-month-old p16-Cre/R26-mTmG and p16-Cre/R26-DTA mice with the vaccine or saline control before analyzing cells from the peritoneal cavity and liver. First, we confirmed a BNT162b2-induced increase in the expression of p16 in control, but not in DTA mice (Figure 3E). We also found increased expression of the regulatory factors *Il-10* and *Arg1* in the peritoneum and liver. In contrast, in DTA mice, *ARG1* expression was diminished, while *IL10* expression was completely abolished (Figure 3E).

### BNT162b2-induced p16^High^ cells protects against severe inflammation and severe tissue damage

Our data indicated that p16^High^ immune subsets could play a role in establishing an immunosuppressive and tissue remodeling environment after treatment with BNT162b2, which in turn could be linked to establishment of disease tolerance, a poorly understood mechanism that attenuates the negative impact of the infection and tissue damage on host fitness^24^. Next sought functional confirmation of these findings. For this purpose, we evaluated the ability of BNT162b2 to acutely protect against severe inflammation and tissue damage under different *in vivo* conditions via p16^High^ cells. Two-month-old control and p16-Cre/R26-DTA mice were primed with BNT162b2 or saline control and 5 days later, the mice were treated intraperitoneal with a high dose of *Escherichia coli* lipopolysaccharide (LPS) to induce sepsis. Sepsis is a life-threatening organ dysfunction affecting more than 48.9 million people worldwide per year caused by a deregulated host response to infections that could be used to robustly evaluate the level of disease tolerance in any particular context^44^. We found that 75% of the saline-treated mice died within 48h of sepsis induction, while only 10% of the BNT162b2-treated mice died. At the termination of the experiment on day 6, the survival rate was 10% and 80% for saline and BNT162b2-treated mice, respectively. Genetic ablation of p16^High^ immune cells resulted in a significant reversal of the protective effect of the vaccine, with only 40% of p16-Cre/R26-DTA mice surviving on day 6 (Figure 3F). Thus, our results indicate that the BNT162b2 vaccine rapidly established disease tolerance by protecting against a high dose of LPS, with p16^High^ cells contributing substantially to this phenomenon.

Encouraged by these results, we decide to extend our investigations to the role of BNT162b2-induced p16^High^ cells in establishing disease tolerance against tissue damage. For this purpose, we delivered a lethal dose of ionizing irradiation (IR). First, we investigated the ability of the BNT162b2 vaccine to increase the number of p16^High^ cells in the intestine as the primary site of tissue damage after this dose of irradiation. While we found only a small but still significant increase in the number of p16^High^ cells in the intestine at day 5 after vaccine treatment, we speculated that a previously identified strong increase in the number of p16^High^ immune cells in the peritoneum (Figure 3A) and liver could restrict intestinal tissue deterioration after irradiation. And this, in turn, could positively contribute to animal survival after an IR. To investigate this further, we irradiated mice with 8 Gy on day 5 after treatment with saline or BNT162b2 and monitored their survival. All saline-treated mice died by day 15, while 40% of the vaccinated control mice were alive and this effect was largely dependent on the presence of p16^High^ cells (Figure 3G). Thus, the BNT162b2 vaccine-induced disease tolerance has a rapid and noticeable radioprotective effect that potentially could be improved by strategies that increase the presence of p16^High^ immune cells in the intestine. The BNT162b2mRNA COVID-19 vaccine has been proven to be very effective in the induction of neutralizing antibodies against a spike protein of the SARS-CoV-2 virus; however, the antibody production starts around day 14 after the priming dose of vaccine both in humans and mice, reaching high levels only by day 21 or after a booster shot^45^. Furthermore, antigen-specific CD8^+^ T cells and increased levels of interferon-γ were detected in mice only after the second dose of the vaccine^45^. If only neutralizing antibodies are considered, these data strongly suggest that the development of a protective immune response against SARS-CoV-2 needs considerable time. Since we found that TLR7-driven induction of p16^High^ immune subsets had almost immediate protective effect against inflammation and tissue damage, we assessed the ability of the BNT162b2 vaccine to promote a rapid and early disease tolerance against lethal inflammation caused by SARS-CoV-2 through induction of p16^High^ immune subsets.

The lungs are the primary site of tissue damage in response to severe SARS-CoV-2 infection. As such, we explored the ability of intraperitoneal injection of BNT162b2 to induce accumulation of p16^High^ immune cell subsets in the lungs. We found a strong increase in p16^High^ expressing immune cells at day 2 after vaccine treatment and this effect was sustained beyond day 15 (Figure 3H). A more detailed analysis of T cells revealed a significant accumulation of Tregs and PD-L1^+^ cells among the p16^High^ T cell populations in the lung (Figure 3I). We further found a strong p16^High^ cell-dependent accumulation of regulatory *Il10* and *Arg1* in the lungs after the treatment with BNT162b2. Together, our data suggest that the BNT162b2 vaccine induces a strong and rapid accumulation of p16^High^ regulatory and immunosuppressive subsets in the lung. This, in turn, could have an immediate tissue protective effect against lethal inflammation. To verify this speculation, we employed a model of viral infection after a lethal dose of a mouse-adapted strain of SARS-CoV-2, MA10^46^. It has been shown that hypercytokinemia (also known as cytokines storm) can, if not contained in a timely manner, eventually lead to Multiple Organ Dysfunction Syndrome (MODS), which is accompanied by rapid weight loss and the eventual death of mice as rapidly as day 2 after infection^47^. Therefore, we analyzed the ability of the BNT162b2 vaccine to provide rapid protection against a lethal dose of SARS-CoV-2 through induction of p16^High^ cells and thus, independently of antibody production. For this purpose, a group of mice was pre-treated with BNT162b2 or PBS (mock) and on day 3, the mice were inoculated intratracheally with a lethal dose of the mouse-adapted SARS-CoV-2 strain MA10. In accordance with previous reports, mice started to lose weight at day 2 after inoculation and this dynamic continued in both groups until day 4, when a rapid recovery began in the BNT162b2-treated mice, while the mock-treated animals continued to lose weight. Analysis of the probability of survival showed a strong protection in the BNT162b2-treated group at 100% survival, while only 30% of the mock-treated mice were alive on day 14 (Figure 3J). Further analysis of lung cytokine gene expression showed a decreased IL6 and *INFb* mRNA expression on day 3 in the BNT162b2-treated group compared to the levels in control mice. Thus, our results provide evidence that mRNA vaccines can almost immediate induce disease tolerance against severe SARS-CoV-2 infection.

### Both TLR7 and STING pathways are required for induction of p16^High^ immune subsets

Next we turn to the analysis of the potential mechanism(s) of induction of p16^High^ immune subsets in response to BNT162b2. It has been previously shown that that immunization with BNT162b2 stimulated potent antibody and antigen-specific T cell responses, as well as strikingly enhanced innate responses after secondary immunization. This mechanism of innate and adaptive immunity to BNT162b2 appeared to be TLR5 and 7- and STING-independent while relied on activation of MDA5^45,48^. To systematically address the role of TLR7, STING and MDA5 pathways in response to BNT162b2 next we used selective inhibitors and knockout mice to dissect the mechanism of p16^High^ immune subset induction. We found that inhibition of TLR7 and STING (with M5049 and H151 respectively) as well as deletion of MDA5 in mice significantly reduced the expression of Isg15 mRNA after treatment with BNT162b2, which was consistent with their role in induction of innate and adaptive immunity to BNT162b2^45,48^. We further found that inhibition of both TLR7 and STING reduced the expression of p16 mRNA (Figure 4A, left panel) after BNT162b2, while surprisingly, MDA5^-/-^ mice showed a significant increase in the basal level of the p16 mRNA expression (Figure 4A, right panel). Because of that, in the first instance we focused on TLR7- and STING-dependent pathways. The use of TLR7 and STING inhibitors blocked the effect of BNT162b2 by reducing the percentage of p16^High^ immune subsets including specifically in Tregs as well as PD1 and PD-L1-positive T cells in analyzed tissues (Figure 4). Next we evaluated an activated and phosphorylated at S366 form of STING in p16^High^ cells. We found that a significant fraction of p16^High^ cells was positive for p-STING with most noticeable co-staining in F4/80^+^ macrophages (Figure 4C&D right panel and upper pie chart). Similarly, F4/80 p16^High^ p-STING^+^ positive cells represented the most significant fraction among CD45^+^ p-STING^+^ cells (Figure 4C&D bottom pie chart). Altogether, our results show that activation of STING after treatment with BNT162b2 coincides with induction of p16^High^ immune subsets.

**Figure 4.**
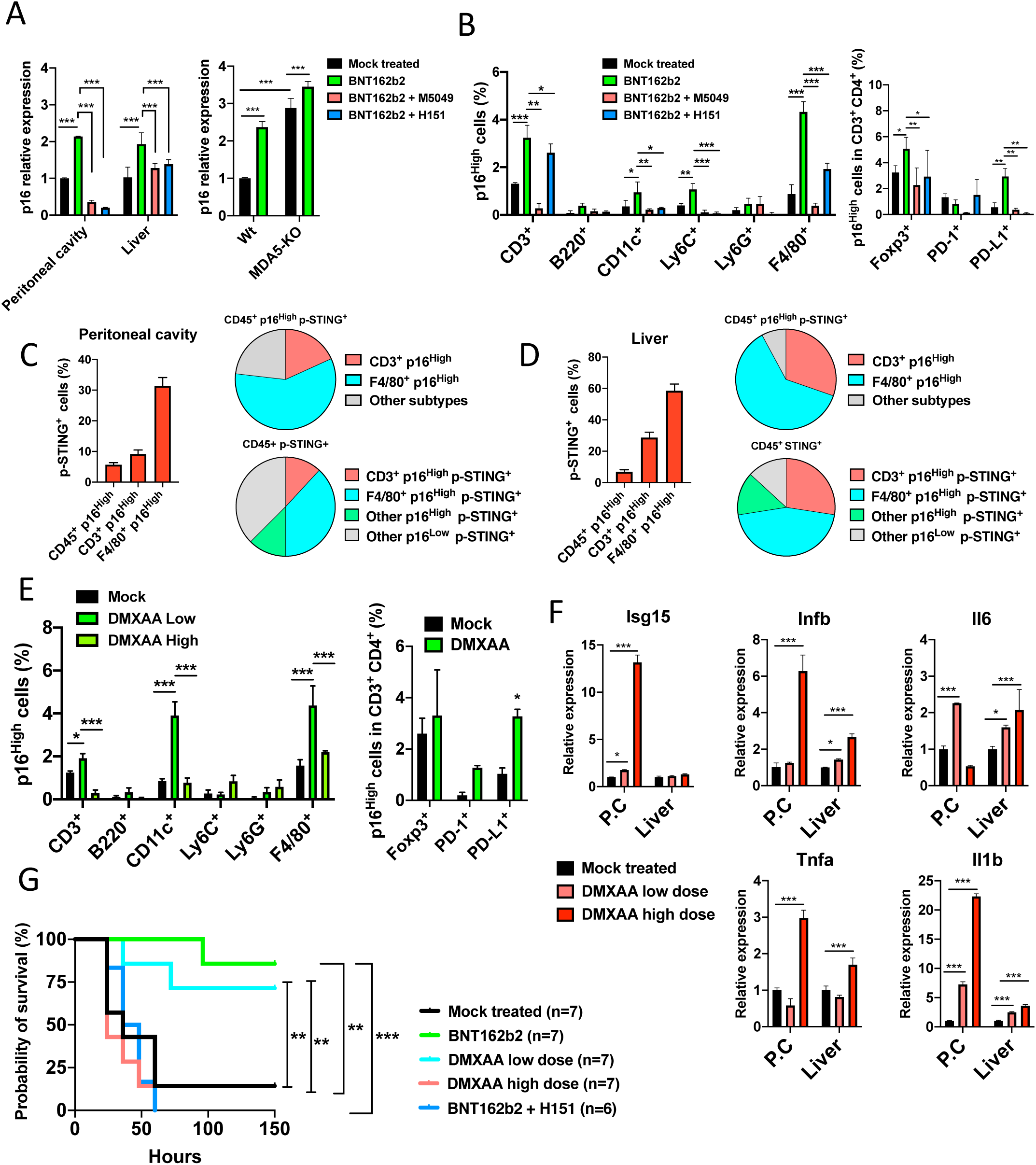
The BNT162b2 mRNA COVID-19 vaccine induces disease tolerance by activation of TLR7 and tonic STING response. **A.** p16 expression is dependent of TLR7 and STING. 2-3 months wilt type mice were treated with the BNT162b2 vaccine. One day before, in the same day, and day after treatment with BNT162b2, some animals were treated I.P with TLR7 inhibitor (M5049, 1 mg/kg) or STING inhibitor (H151, 10 mg/kg). p16 mRNA expression was determined after treatment in peritoneal cells and liver (left panel) by qPCR. 2-3 months MDA5^+/+^ (wild type) and MDA5^-/-^ (MDA5-KO) mice were treated with the BNT162b2 vaccine. After 5 days of treatment p16 mRNA expression was determined after treatment in liver (right panel) by qPCR. Data are mean +/- S.D. Difference between groups were analyzed using ANOVA test and Tukey post hoc test. ***p<0,001. *n = 3* biologically independent samples. **B.** 2-3 months p16-Cre/R26-mTmG mice were treated with the BNT162b2 vaccine. One day before, in the same day, and day after treatment with BNT162b2, some animals were treated I.P with TLR7 inhibitor (M5049, 1 mg/kg) or STING inhibitor (H151, 10 mg/kg). 5 days after BNT162b2 treatment, animals were euthanized and single cells suspensions were analyzed by flow cytometry to determine the percentage of p16^High^ cells on different immune subsets (left panel) and inside of Foxp3, PD-1 and PD-L1 (CD3^+^ CD4^+^) populations (right panel). *n = 3* biologically independent samples. Data are mean +/- S.D. Differences between groups were analyzed using ANOVA test and Tukey post hoc test. *p<0,05; **p<0,01 and ***p<0,001. **C**-**D.** 4-6 months p16-Cre/R26-mTmG mice were treated with the BNT162b2 vaccine. 5 days. 5 days after BNT162b2 treatment, animals were euthanized and single cells suspensions from peritoneal cavity (**C**) and liver (**D**) were analyzed by flow cytometry to determine the percentage of activated STING (p-STING) (phosphorylated at Ser366) in CD45^+^ p16^High^, F4/80^+^ p16^High^, and CD3^+^ p16^High^ populations (left panel). The percentage covered of F4/80^+^ p16^High^, and CD3^+^ p16^High^ in the total CD45^+^ p16^High^ p-STING^+^ is shown in the upper pie chart. The p-STING^+^ p16^High^ F4/80^+^ and CD3^+^ abundance and distribution were determined in the total CD45+ p-STING^+^ and is show in the lower pie chart. *n = 3* biologically independent samples. Data are mean +/- S.D. **E.** STING activation induces p16^High^ subsets. 2-3 months p16-Cre/R26-mTmG mice were treated I.P with the STING agonist DMXAA at low or high dose and 5 days after treatment, animals were euthanized and cells suspensions from peritoneal cavity were analyzed by flow cytometry to determine the percentage of p16^High^ cells on different immune subsets (left panel). Differences between groups were analyzed using ANOVA test plus Tukey post hoc test. For the low dose condition, percentage of p16^High^ cells in Foxp3, PD-1 and PD-L1 (CD3^+^ CD4^+^) populations (right panel) was determined. Data are mean +/- S.D. Differences between groups were analyzed using ANOVA test plus Dunnett post hoc test. *p<0,05; **p<0,01 and ***p<0,001. *n = 3* biologically independent samples. **F.** 2-3 months wild type mice were treated I.P with the STING agonist DMXAA (10 mg/kg) one time (low dose) or 2 consecutive days (high dose). 5 days after first treatment, animals were euthanized and RNA from peritoneal cells (P.C), and liver was isolated. RNA was analyzed by qPCR to determine the level of expression of inflammatory genes. Data are mean +/- S.D. Independent repeats *n = 3.* Differences between groups were analyzed using ANOVA test plus Dunnett post hoc test. *p<0,05; and ***p<0,001. **G.** Tonic STING activation promotes disease tolerance and tissue protection against severe inflammation. Wild type mice were either treated with saline (Mock); BNT162b2 vaccine (5 µg by mouse); BNT162b2 vaccine plus H151 (10 mg/kg, 3 consecutive days starting one day before BNT162b2 treatment); and DMXAA (10 mg/kg) for one (low dose) or two consecutive days subcutaneously (high dose). After 5 days, animals were injected with LPS O55:B5 30 mg/kg. Animals were euthanized immediately once they reached the limit point of physical deterioration. Difference between groups was analyzed using Gehan-Breslow-Wilcoxon test. **p<0,01 and ***p<0,001. *n* used for each condition is showed in the plot.

Next we tested and found that a direct activation of STING with the specific agonist DMXAA at low doses was sufficient to increase the number of p16^High^ immune subsets including within Tregs as well as PD1- and PD-L1-positive T cells in different tissues (Figure 4E). In contrast, high doses of DMXAA resulted in significantly reduced numbers of p16^High^ immune subsets both in the peritoneum and liver (Figure 4E). While STING activation at low doses was important in induction of p16^High^ immune subsets, its activation has been broadly implicated in the regulation of numerous pro-inflammatory and pro-apoptotic pathways which is in part due to its capacity to induce TNFα^49,50^. This in turn could result in increased apoptosis of immune cells and explain lower numbers of p16^High^ immune subsets after high doses of DMXAA. Consistent with these predictions, we found that an increase dose of the STING agonist, in contrast to low doses, strongly induced the expression of pro-inflammatory factors including *Il1b, IL6* and importantly *TNF*α (Figure 4F).

Next we analyzed the role of a direct STING activation on protecting mice from LPS-induced sepsis. We found that while low doses of the STING activator DMXAA phenocopied the effect of BNT162b2 on protecting mice from LPS-induced severe inflammation, significant STING activation or inhibition after BNT162b2 no longer had a protective effect (Figure 4G). Thus, low levels of STING activation could be beneficial while high – detrimental due to induction of excessive inflammation, in protecting mice from LPS-induced sepsis.

### Nnmt is a key molecule controlling p16^High^ immune subsets

To gain further insights into the potential mechanism(s) of p16 induction in immune cells, we evaluated genes that were found to be differentially expressed in p16^High^ versus p16^Low^ F4/80^+^ peritoneal macrophages based on RNA-Seq. We found that some of the most upregulated genes included nicotinamide N-methyltransferase (Nnmt) (Figure 5A), which regulates numerous physiologically-relevant processes by controlling the availability of S-adenosyl methionine (SAM), the main substrate for all types of methylation. Furthermore, we recently identified an important role of Nnmt in the induction of p16^High^ fibroblasts during iPSC reprogramming^51^, providing further credence to the role of Nnmt as a potentially important molecule relevant to a program controlling p16 activation.

**Figure 5.**
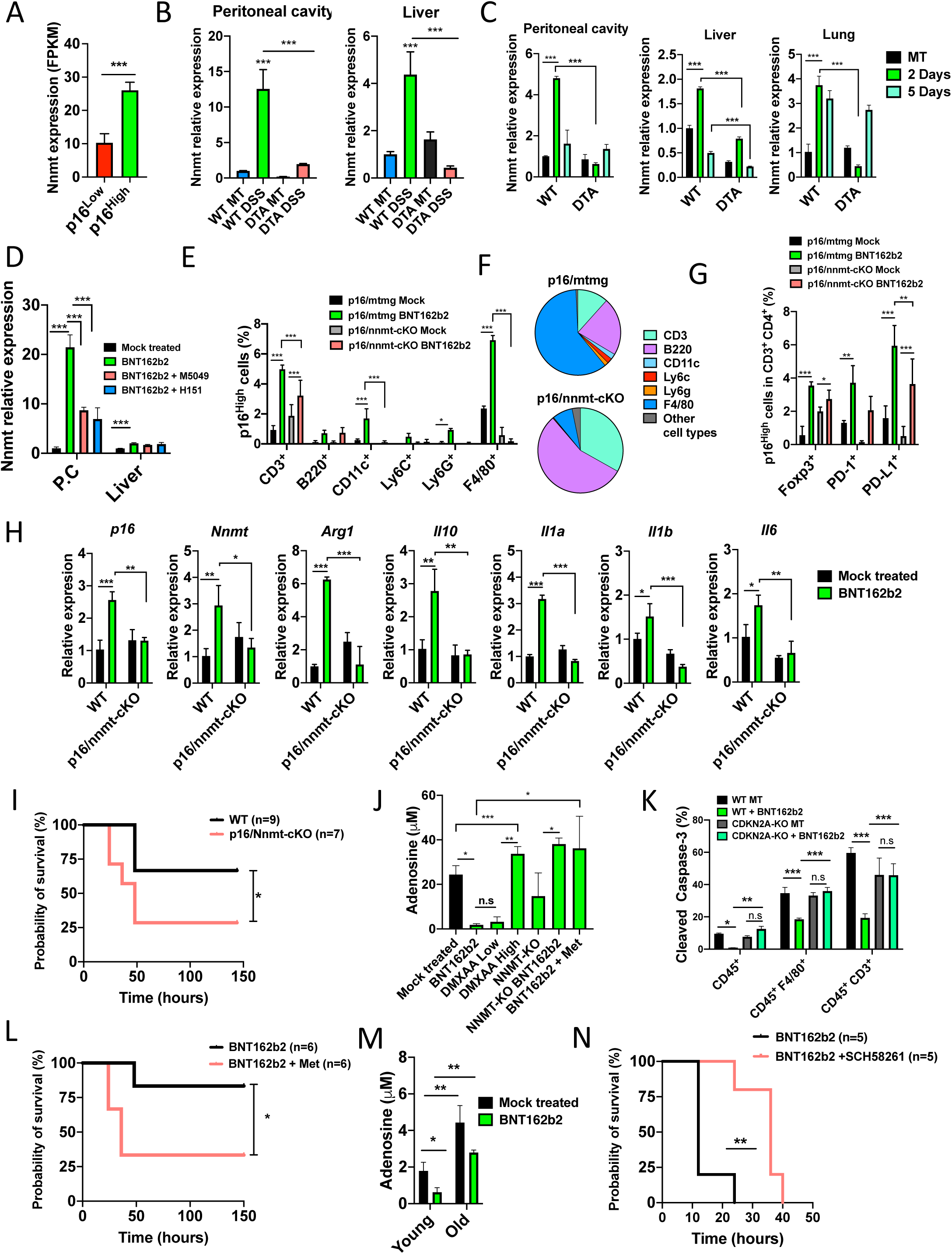
Nnmt expression and low adenosine are key conditions to induce p16^High^ immune subsets and disease tolerance. **A.** The Nnmt gene is overexpressed in p16^High^ cells. RNA-seq data reveals overexpression of Nnmt gene in p16^High^ cells. Fragments Per Kilobase of transcript sequence per Millions base pairs sequenced (FPKM) from p16^High^ and p16^low^ populations were analyzed by t-test. *n = 3* biologically independent samples. Data are mean ± SD. ***p<0,001. **B.** The Nnmt gene is induced after DSS treatment. 2-3 month-old control and p16-Cre/R26-DTA (DTA) were treated with 2.5% DSS for 7 days and RNA from peritoneal cells and liver was isolated on day 8. RNA was analyzed by a quantitative SYBR-green based PCR (qPCR) to determine the level of expression Nnmt mRNA. Independent repeats *n = 3.* Data are mean +/- S.D. Difference between groups was analyzed using ANOVA test and Tukey post hoc test. ***p<0,001. **C.** BNT162b2 vaccine-induced Nnmt expression depends of p16^High^ cells. 2-3 month-old control and p16-Cre/R26-DTA (DTA) were treated with 5 µg of BNT162b2 vaccine. After 2 and 5 days, RNA from peritoneal cells, liver and lungs was isolated and analyzed by qPCR to determine the level of expression of Nnmt mRNA. Independent repeats *n = 3.* Data are mean +/- SD. Difference between groups was analyzed using ANOVA test and Tukey post hoc test. *p<0,05; **p<0,01 and ***p<0,001. **D.** Nnmt expression is dependent of TLR7 and STING. 2-3 months wilt type mice were treated with the BNT162b2 vaccine. One day before, in the same day, and day after treatment with BNT162b2 some animals were treated I.P with TLR7 inhibitor (M5049, 1 mg/kg) or STING inhibitor (H151, 10 mg/kg). Nnmt mRNA expression was determined after treatment in peritoneal cells (P.C) and liver (left panel) by qPCR. Independent repeats *n = 3.* Data are mean +/- S.D. Difference between groups were analyzed using ANOVA test and Tukey post hoc test. ***p<0,001 **E.** Nnmt is required to maintain p16^High^ immune cells. 2-3 month-old p16-Cre/R26-mTmG and p16-Cre/Nnmt-cKO (Nnmt^-/-^ conditional to the expression of p16) mice were treated intraperitoneal with 5 µg of BNT162b2 vaccine. After 5 days single cell suspensions from peritoneal cavity were stained with conjugated antibodies against different immune cell types. Cells suspensions were analyzed by flow cytometry and the percentage of p16^High^ cells in each population was determined. Independent repeats *n = 3.* Data are mean ± SD. Statistical significance were determined using ANOVA plus Tuckey post hoc test. ***p<0,001. **F.** Distribution and abundance of total p16^High^ cells from peritoneal cavity in p16-Cre/R26-mTmG and p16/Nnmt-cKO after BNT162b2 vaccine treatment are shown. **G.** Percentage of p16^High^ positive cells in different T cell populations was determined during experiment show in **E**. Data are mean ± SD. Statistical significance were determined using ANOVA plus Tukey post hoc test. ***p<0,001. **H.** Mice with selective inactivation of the Nnmt gene in p16^High^ cells cannot mount full immune response. 2-3 month-old control and p16/Nnmt-cKO mice were treated with 5 µg of BNT162b2 vaccine. 5 days after treatment, RNA from peritoneal cells was isolated and analyzed by quantitative PCR to determine the level of expression of Nnmt, p16 and inflammatory genes. Data are mean +/- S.D. Independent repeats *n = 3.* Differences between groups were analyzed using ANOVA test and Tukey post hoc test. *p<0,05; **p<0,01 and ***p<0,001. **I.** Nnmt is necessary to protect against acute inflammation. 2-3 months wild type and p16/Nnmt-cKO mice were treated with 5 µg of BNT162b2 vaccine. After 5 days, animals were injected I.P with LPS O55:B5 40 mg/kg. Animals were euthanized immediately once they reached the limit point of physical deterioration. Difference between groups was analyzed using Gehan-Breslow-Wilcoxon test. *p<0,05. *n* used for each condition is showed in the plot. **J.** Activation of p16^High^ program lower adenosine level. 2-3 months wild type mice were treated I.P with saline (Mock), BNT162b2 vaccine, one (low) or two doses (high) of DMXAA (10 mg/kg), and BNT162b2 (µg by mouse) plus orally supplemented L-methionine daily during 5 days (200 mg/kg). Additionally, 2-3 months p16/Nnmt-cKO mice (written as Nnmt-KO in graph label) were treated with saline or BNT162b2 vaccine. After treatment, peritoneal cells were collected and lysed, levels of adenosine were determined in the supernatant. Data are mean +/- S.D. Differences between groups were analyzed using ANOVA test and Tukey post hoc test. *p<0,05; **p<0,01 and ***p<0,001. **K.** 3-4 months CDKN2A^-/-^ (CDKN2A-KO) and CDKN2A^+/+^ (wild type) mice were either treated with saline (Mock) or BNT162b2 vaccine (5 µg by mouse). Single cell suspensions from peritoneal cavity were analyzed by flow cytometry to determine the percentage of cleaved-caspase-3 in CD45, F4/80 and CD3 populations. Data are mean +/- S.D. Differences between groups were analyzed using ANOVA test and Tukey post hoc test. *p<0,05; **p<0,01 and ***p<0,001. **L.** High adenosine levels reduce BNT162b2 effect on survival. 2-3 months wild type mice were treated with 5 µg of BNT162b2 vaccine, or BNT162b2 plus daily supplementation with L-methionine (200 mg/kg). After 5 days, animals were injected I.P with LPS O55:B5 30 mg/kg. Animals were euthanized immediately once they reached the limit point of physical deterioration. Difference between groups was analyzed using Gehan-Breslow-Wilcoxon test. *p<0,05. *n* used for each condition is showed in the plot. **M.** Adenosine level is increase during aging. 3-4 months and 20 months-old wild type mice were treated with saline or BNT162b2 vaccine (5 µg by mouse). 5 days after animals were euthanized and levels of adenosine were determined in CD45^+^ liver cells. Independent repeats *n = 3.* Data are mean +/- S.D. Differences between groups were analyzed using ANOVA test and Tukey post hoc test. *p<0,05 and **p<0,01. **N.** Blocking adenosine receptor A_2A_ increase survival during aging. 2 year-old wild type mice were treated with 5 µg of BNT162b2 vaccine, or BNT162b2 plus adenosine receptor A_2A_ inhibitor (SCH58261, twice by day, 3 mg/kg) starting one day before LPS treatment. 5 days after BNT162b2 treatment, animals were injected I.P with LPS O55:B5 20 mg/kg. Animals were euthanized immediately once they reached the limit point of physical deterioration. Difference between groups was analyzed using Gehan-Breslow-Wilcoxon test. **p<0,01. *n* used for each condition is showed in the plot.

First, we determined the level of Nnmt after treatment with DSS *in vivo* and found its significant induction in the peritoneum and liver in wild-type mice, but not in p16-Cre/R26-DTA mice (Figure 5B). Similar results with upregulation of Nnmt mRNA were obtained after treatment with BNT162b2 in different tissues (Figure 5C) while this effect was dependent on both TLR7 and STING (Figure 5D).

To gain additional insight into the role of NNMT in controlling p16^High^ immune subsets *in vivo*, and due to low bioavailability and activity of current NNMT chemical inhibitors, we next generated Nnmt conditional knockout (Nnmt-cKO) mice. In these mice, the first exon of the *Nnmt* gene was flanked by LoxP sites to produce an inactive gene after excision with the Cre recombinase. Subsequently, we crossed Nnmt-cKO mice with mice expressing Cre under the control of the p16 promoter (p16-Cre). The resulting animals in which the removal of Nnmt was induced specifically in p16^High^ cells were used for further experimentation. We found that the ability of BNT162b2 to induce p16^High^ subsets was greatly diminished in different tissues in the p16-Cre/Nnmt-cKO mice (Figure 5E&F) including in Tregs, PD1- and PD-L1-positive T cells (Figure 5G). This correlated with the lack of p16, Nnmt, IL-10 and Arg1 mRNA induction after BNT162b2 treatment in p16/Nnmt-cKO mice (Figure 5H), indicating that Nnmt activation is a critical mechanism of p16 induction either directly or indirectly. In addition, the cells from the peritoneal cavity of p16-Cre/Nnmt-cKO mice were unable to mount a full inflammatory response (Figure 5H).

As conditional deletion of Nnmt produced a phenotype that resembled the *in vivo* features of p16-Cre/R26-DTA mice, including the reduced number of p16^High^ immune cells and attenuated inflammatory response (Figure 5H), we hypothesized that these mice would also have a reduced survival in response to LPS-induced sepsis. To test this hypothesis, we treated wild type and p16-Cre/Nnmt-cKO mice with BNT162b2 and after 5 days, injected a high dose of LPS. We found that the ability of BNT162b2-treated mice to survive LPS-induced sepsis was greatly reduced in p16-Cre/Nnmt-cKO mice (Figure 5I), resembling our previous observations in p16-Cre/R26-DTA mice (Figure 3F). Overall, our data support a critical role of Nnmt in inducing p16^High^ immune cell subsets and protection from LPS-induced sepsis.

### p16^High^ immune cells protect from severe inflammation by reducing the levels of adenosine

Overexpression of the Nnmt gene leads to reduced levels of S-Adenosyl Methionine (SAM) in p16^High^ cells^51^, which could result in attenuated adenosine concentrations after SAH hydrolysis. Adenosine is an important regulator of the immune system, with strong immunosuppressive properties^52,53^. In turn, adenosine build up could derail a balanced immune response leading to reduced survival of animals during severe inflammation. Consistent with the role of NNMT in controlling SAM levels and potentially adenosine, the BNT162b2 treatment significantly reduced adenosine concentrations in wild-type but not in p16-Cre/Nnmt-cKO mice (Figure 5J). Same effect was observed with low but not high STING activation. Importantly, supplementation with L-methionine, a main precursor of SAM, fully reversed the effect of BNT162b2 on reducing the levels of adenosine (Figure 5J). Thus, BNT162b2 lowers adenosine concentrations via NNMT and reduction of SAM levels.

While the level of adenosine could be controlled via SAM, one of the major sources of adenosine, and specifically extracellular adenosine, is the ATP. ATP is released to the environment from apoptotic cells in damaged and inflamed tissues^54^, including regulatory T cells^55^. In turn, inhibition of the cell cycle, including by upregulation of p16, has been reported to have anti-apoptotic properties ^56^. In addition, p16^High^ cells commonly express the high level of anti-apoptotic protein Bcl2^57^. Because of that, we hypothesize that p16 upregulation may protect immune cells from apoptosis which in turn could lead to reduction of extracellular adenosine. To verify that, we treated 2-month-old wild type and Cdkn2a knockout (Cdkn2a-KO) mice with BNT162b2 and 5 days later exposed them to sub-lethal doses of LPS. Analysis of cleaved-caspase 3-positive cells 12h after LPS treatment showed that BNT162b2 effectively reduced the activation of apoptosis in CD45, F4/80 and CD3 populations from wild type but not Cdkn2a-KO mice (Figure 5K). Thus, upregulation of p16 could be critical in counterbalancing apoptosis in immune cells during severe inflammation.

Next, we check whether controlling adenosine levels or adenosine signaling could have an impact on protecting mice from LPS-induced sepsis. For that, we treated 2-month-old animals with BNT162b2 with and without daily supplementation of L-methionine. We found that L-methionine-supplemented animals showed a significantly lower survival rate in a model of LPS-induced sepsis (Figure 5L). Since we found that maintaining low adenosine levels are critical for development of BNT162b2-induced protection from LPS-induced sepsis, next we turned to old (24 months) animals to evaluate their response to LPS. It is well established that during aging, adenosine levels are significantly increased most likely due to a much higher rate of cell death in aged tissues^58,59^. We further confirmed an increase in adenosine levels in different tissues of old (24 months) versus young (2 months) animals (Figure 5M). Interestingly, while BNT162b2 was able to lower adenosine in old animals, this reduction has never reached the level observed in young mice (Figure 5M). Consistent with inability to significantly downregulate adenosine levels after the treatment with BNT162b2 in old mice, we found that BNT162b2 completely failed to protect 24-month-old mice from LPS-induced sepsis (Figure 5M). In contrast, a simultaneous treatment with the adenosine receptor A_2A_ inhibitor SCH58261 was sufficient to significantly improve the protective effect of BNT162b2 from LPS-induced sepsis in old mice (Figure 5N). Thus, low adenosine levels are essential for BNT162b2-induced protection from LPS-induced sepsis both in young and old animals.

### Deficiency of MDA5 promotes healthspan

Our findings show that the presence of p16^High^ immune subsets is indispensable for animal survival in response to multiple lethal conditions such as LPS-induced sepsis, acute COVID-19 infection, as well as Ionizing irradiation (Figures 3F,G,K). In turn, this could argue that maintaining an increased number of p16^High^ immune cells subsets from early age and thus before any tissue damage or inflammation could potentially provide benefits in extending healthspan. As we previously found a higher basal level of p16 expression in young MDA5-KO (Ifih1^-/-^) mice (Figure 4A) we wanted to explore whether indeed such an increase could provide bases for improved healthspan. Since deletion of MDA5 could potentially induce STING as a feedback mechanism due to lack of sensing of endogenous double-stranded RNAs including from repetitive and antisense sequences^60^, next we seek to verify whether higher p16 expression in the tissues of MDA5-KO mice was indeed dependent on STING. Our analysis of an activated and phosphorylated form of STING in different immune populations (CD45, CD3, F4/80) showed their increase in MDA5-deficient cells which was significantly attenuated in the presence of the STING inhibitor, H151 (Figure 6A). p16 expression in MDA5 KO mice was reduced after H151 treatment (Figure 6B). Next, we moved to analyze aged MDA5 KO mice. The level of adenosine rises with age (Figure 5M and Figure 6C) while maintaining it low is a pre-requisite for increased organismal survival in response to multiple conditions as we describe here. Consistent with upregulation of p16 mRNA, our analysis of adenosine concentrations in MDA5 KO cells showed its significantly reduced levels (Figure 6C). Indeed, these levels were close to the ones observed in wild type mice after treatment with BNT162b2 suggesting that MDA5 KO mice are potentially in a continuously primed state of heighten resistance to different tissue damaging conditions and thus their tissue deterioration with aging could occur at a slower pace. To test that, we analyzed 24-month-old wild type and MDA5-KO littermates mice and found a significant reduction in the level of expression of pro-inflammatory cytokines IL-1 (α, but not β), IL-6, TNF-α, and Cxcl13, while an increase in the expression of anti-inflammatory IL-10 (Figure 6D). Next we checked whether such a delay in developing of tissue inflammation with aging correlated with improved histological characteristics of different tissues. Among such characteristics, we checked the level of tissue fibrosis and blood vascularization. The latter is particularly critical since it is significantly reduced with aging^51^ while its improvement in VEGF transgenic mice is sufficient to extend health- and life-span^61^. We found that the level of αSMA, a marker of fibrosis, was significantly reduced while the presence of Cd31-positive cells, a marker of vascular endothelium, was strongly increased in different tissues from MDA5-KO when compared to wild type littermates (Figure 6F). To test further whether observed changes in MDA5 KO mice had functional and physiological significance, next we determined muscle strength based on a grip test and found it significant improvement in MDA5 KO mice when compared to wild type littermates (Figure 6G). Encouraged by these results, we evaluated the general physical state of animals based on a frailty score. We found that aged MDA5-KO in comparison to wild type littermates showed a significantly reduced frailty index (Figure 6H). Finally, both wild type and MDA5-KO littermates have been followed for 26 months to evaluate the median survival. We found that both male and female MDA5 KO mice showed a significantly improved median survival with aging (Figure 6I) further supporting a model of targeting MDA5 for healthspan extension.

**Figure 6.**
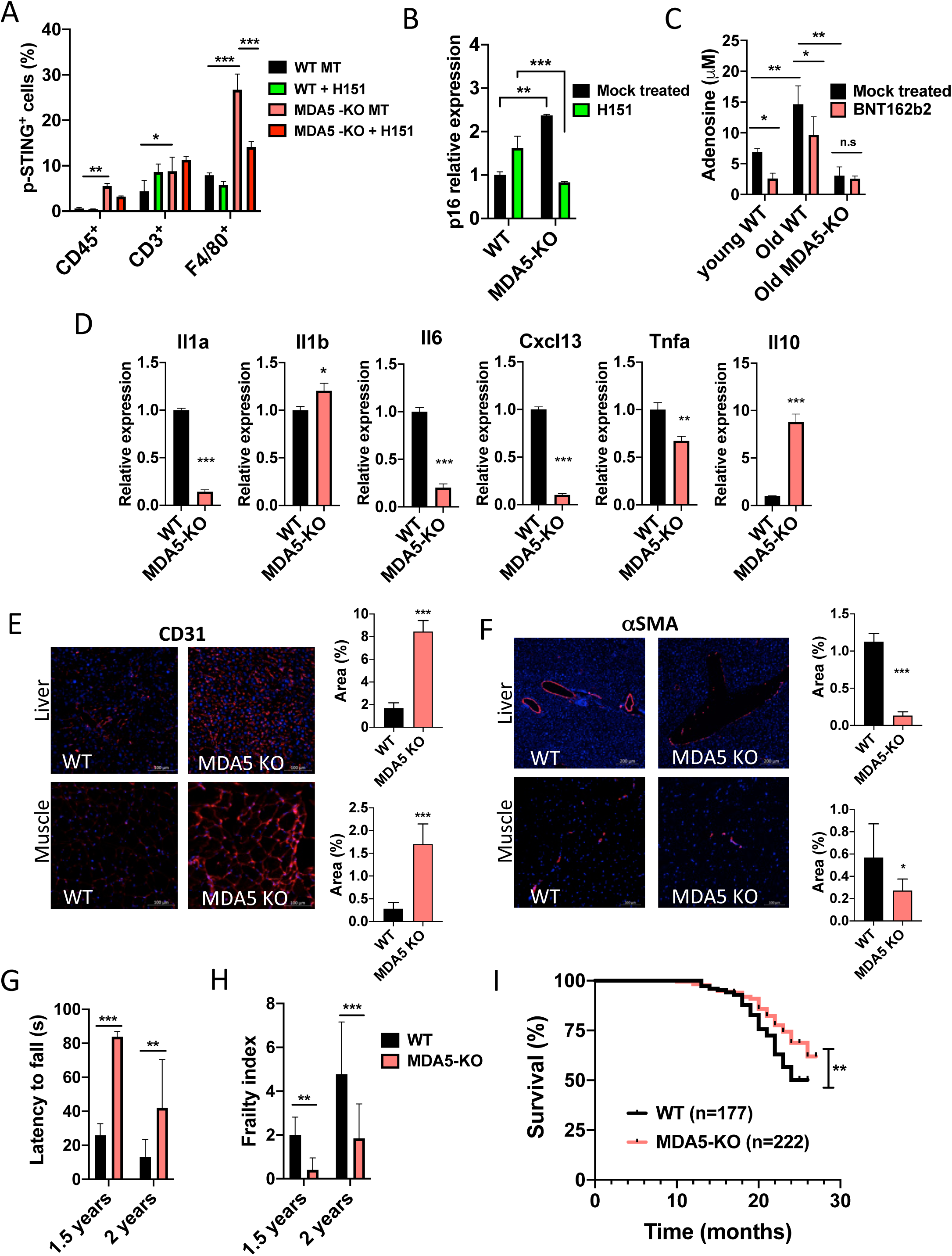
MDA5 downregulation promotes increased physiological fitness during natural aging. **A.** Activated STING is increased in young MDA5-KO mice. 3-4 months Ifih1^+/+^ (wild type) and Ifih1^-/-^ (MDA5-KO) littermates were treated with saline (Mock) or STING inhibitor (H151, 10 mg/kg). 48 h after, animals were euthanized and single cells suspensions from liver were analyzed by flow cytometry. Percentage of p-STING-Ser366 (and p-positive cells was determined in CD45, F4/80 and CD3 populations. *n* used for each condition is showed in the plot.Data are mean +/- S.D. Differences between groups were analyzed using ANOVA test and Tukey post hoc test. *p<0,05; **p<0,01 and ***p<0,001. **B.** p16 is overexpressed in MDA5-KO mice. 18-20 months MDA5^+/+^ (wild type) and MDA5^-/-^ (MDA5-KO) littermates were treated with saline or H151 (10 mg/kg). 48 h after, animals were euthanized and mRNA from livers was isolated and analyzed by qPCR. Independent repeats *n = 3.* Data are mean +/- S.D. Differences between groups were analyzed using ANOVA test and Tukey post hoc test. **p<0,01 and ***p<0,001. **C.** MDA5-KO mice exhibit reduced adenosine levels during aging. 3-4 months MDA5^+/+^ (wild type); and 20 months Ifih1^+/+^ (wild type) Ifih1^-/-^ (MDA5-KO) littermates were treated with saline or BNT162b2 vaccine (5 µg by mouse). 5 days after animals were euthanized and levels of adenosine were determined in peritoneal cells. Independent repeats *n = 3.* Data are mean +/- S.D. Differences between groups were analyzed using ANOVA test and Tukey post hoc test. *p<0,05; **p<0,01 and ***p<0,001. **D.** MDA5-KO mice shown less basal inflammation during aging. Liver RNA from 3-4 months Ifih1^+/+^ (wild type) and Ifih1^-/-^ (MDA5-KO) littermates was isolated and analyzed by qPCR. The level of expression of inflammatory genes was determined. Independent repeats *n = 3.* Data are mean +/- S.D. Differences between groups were analyzed using t-test. *p<0,05; **p<0,01 and ***p<0,001. **E.** Liver and muscle samples isolated from 24-month old wild type and MDA5-KO mice, stained for the endothelial marker CD31. Graph at the right represents mean ± S.D of CD31+ area. Area was determined among 3 animals per group. Differences between groups were calculated by unpaired, nonparametric Mann-Whitney test. ***p<0,001. Scale bar – 100 mm. **F.** Immunofluorescent analysis of fibrosis marker aSMA in liver and muscle samples (same as in **E**). Graphs at the right represent mean ± S.D of aSMA area between groups, Area was determined among 3 animals per group. *n = 3* biologically independent samples. Differences between groups were determined by unpaired, nonparametric Mann-Whitney test. *p<0,05; and ***p<0,001. Scale bars: 200 mm for liver, 100 mm for muscle. **G.** Muscle strength in 18 months wild type (*n = 4*) and MDA5-KO (*n = 5*); and 24 months old months wild type (*n = 10*) and MDA5-KO (*n = 15*) littermates mice was determined with a grip test. Data are mean +/- S.D. Differences between groups were analyzed using ANOVA test and Dunnett post hoc test. **p<0,01 and ***p<0,001. **H.** MDA5-KO mice show better physical fitness during natural aging. Frailty was assessed in 18 months wild type (*n = 4*) and MDA5-KO (*n = 5*); and 24 months old months wild type (*n = 10*) and MDA5-KO (*n = 15*) littermates using an clinically relevant index for frailty during aging (see material and methods). Data are mean +/- S.D. Differences between groups were analyzed using ANOVA test and Dunnett post hoc test. **p<0,01 and ***p<0,001. **I.** A group of wild type (*n = 177*) and MDA5-KO (*n = 222*) animals were housed in the same conditions and followed during 25 months to determine survival during natural aging. Difference between groups was analyzed using Gehan-Breslow-Wilcoxon test. **p<0,01.

### The BNT162b2 vaccine induces p16^+^ immune subsets in humans that are significantly reduced in severe COVID-19 patients

To confirm that the BNT161b2 vaccine could induce p16^High^ immune cells in humans, we evaluated peripheral blood before and after standard vaccination with BNT162b2. Seven healthy donors (4 women and 3 men) were sampled prior vaccination, as well as 7 and 30 days later. The mean age of the cohort was 36 years [23; 42]. Most had already received at least three doses of RNA vaccines (either BNT162b2 or mRNA-1273), with the last dose >6 months earlier (according to French recommendations for health care workers), and ≥1 SARS-CoV-2 infection in the past 3 years. The 4^th^ dose of the vaccine (BNT162b2 only) was well-tolerated by all participants and did not induce an inflammatory syndrome (increased IL-1β and IL-6), but produced a significant SARS-CoV2-specific T cell response 1 month later (Supplementary Table 1). Next, we performed RT-PCR analysis of isolated PBMCs and confirmed an increase in the level of p16 mRNA after vaccination (Figure 7A). In the same way, the expression of NNMT, which we found to be upstream of p16 activation in immune cells was also increased after vaccination in humans (Figure 7B). To further validate our results, we performed flow cytometry analysis in freshly sampled whole blood and found a strong increase in the percentage of cells expressing p16 within the CD45^+^ cell populations at 7d which are significantly subsided by 30d after vaccination (Figure 7C&D). All of the analyzed immune cell subtypes, including granulocytes, monocytes, lymphocytes but most importantly Tregs, showed significantly increased expression of p16 protein at 7d (Figure 7D&E). We further observed a strong induction of p16 protein in PD1-positive immune cells after vaccination (Figure 7C&D). These changes accompanied by a significant increase in the level of plasma regulatory IL10 after vaccination (Table 1). Thus, standard vaccination with BNT162b2 in humans transitory induces p16^High^ immune cell subsets, which in turn could be critical to provide a rapid and broad tissue protection from severe inflammation before development of inhibitory antibody.

**Figure 7.**
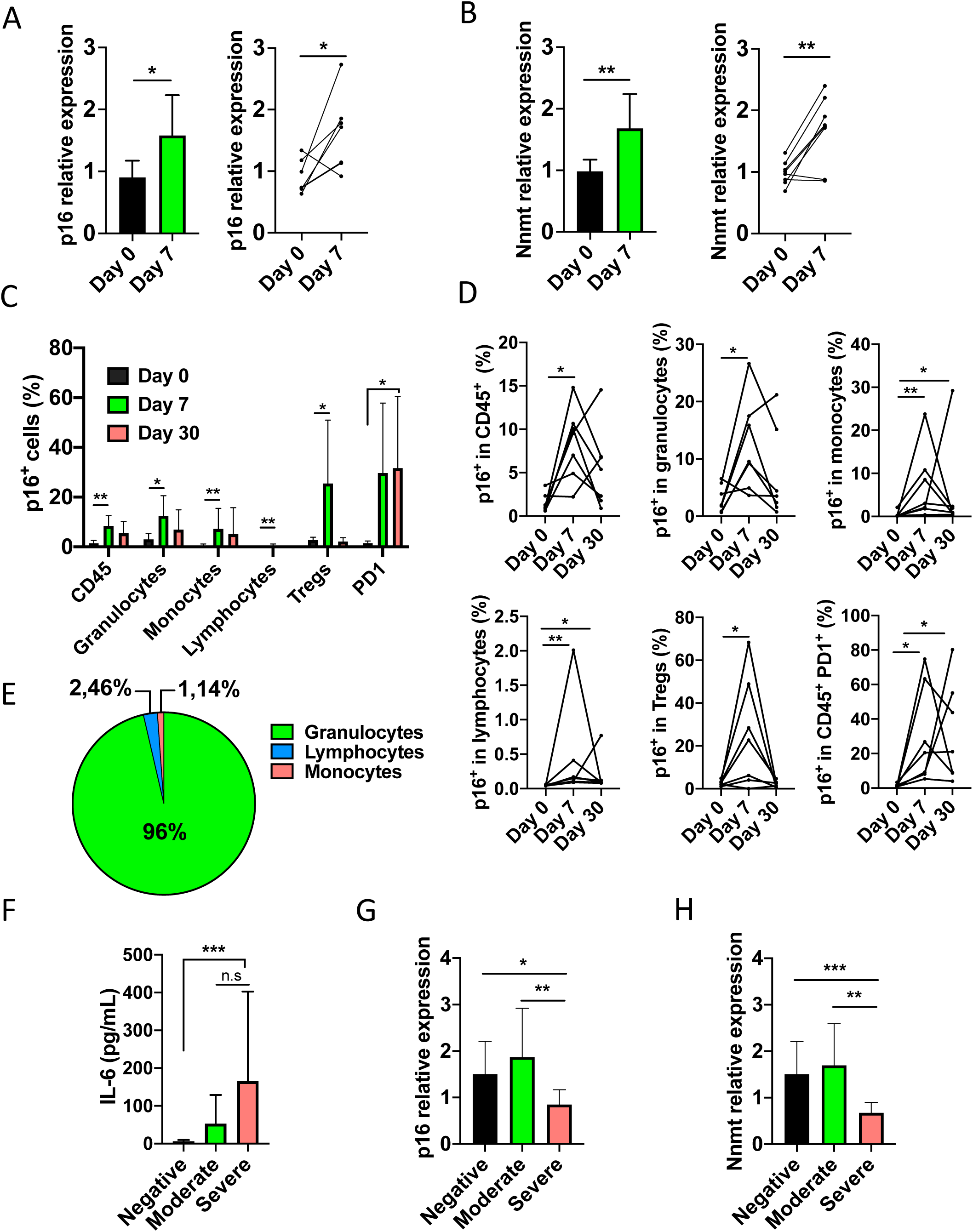
The BNT162b2 vaccine induces p16^+^ immune subsets in humans that are significantly reduced in severe COVID-19 patients. **A-B.** Expression of p16 and NNMT before (Day 0) and after BNT162b2 vaccine treatment (Day 7). RNA were isolated from PBMCs and analyzed by quantitative-PCR. (7 participants were involved in the study). Data were analyzed as a bulk (left panel) and as matched samples (right panel). t-test and paired t-test (right panel) were used to determine the differences between conditions. *p<0,05 and **p<0,01. **C.** Levels of p16^+^ cells in different immune populations before and after BNT162b2 treatment are shown. 7 volunteers (included in the CovImmune 2 cohort) with different range of age were involved in the study. Volunteers were treated intramuscularly with 30 µg of the BNT162b2 vaccine. Blood samples were collected before vaccination and after 7 and 30 days of treatment. Blood samples were analyzed by flow cytometry. Granulocytes, monocytes and lymphocytes populations were localized through size and granularity. CD45^+^, Tregs (CD3^+^ CD4^+^ Foxp3^+^), PD1^+^, and p16^+^ populations were localized with fluorescent conjugated antibodies. Data were analyzed as a bulk. Differences were determined using ANOVA test and Dunnetts test. *p<0,05 and ***p<0,001. **D.** Flow cytometry data were also treated as matched samples. The level of p16^+^ cells inside singles populations are shown in each plot. Paired t-test was used to determine the differences before (Day 0) and after (Day 7 and day 30) the BNT162b2 vaccine treatment for CD45^+^, granulocytes, Tregs and PD1^+^. For monocytes and lymphocytes, Shapiro-Wilk test was used to determine normality of the data. ANOVA plus Dunnett’s test or Friedman plus Dunńs test were used to determine difference before and after. *p<0,05 and **p<0,01. **E.** The total number of cells in CD45^+^, lymphocytes, granulocytes and monocytes, was used to determine the distribution of p16^+^ cells among these populations. The percentage of contribution calculated for each population is showed in the pay graph at day 7. **F.** One cohort (CovImmune 1) of 41 unvaccinated patients and non-COVID volunteers older than 55 years were classified in negative for COVID-19 (*n = 17*), moderate COVID-19 (*n = 6*) and severe COVID-19 (*n = 18*). IL6 concentration in serum was analyzed by ELISA. Data are mean +/- S.D. Differences between groups were analyzed using ANOVA test and Tukey post hoc test. ***p<0,001. **G, H.** One cohort of 41 unvaccinated patients in a older than 55 years were classified in negative for COVID-19 (*n = 17*), moderate COVID-19 (*n = 6*) and severe COVID-19 (*n = 18*). RNA from PBMCs were isolated and analyzed by quantitative-PCR. Level of p16 (**G**) and NNMT (**H**) expression were determined. Differences among groups were determined using ANOVA test and Tukey post hoc test. *p<0,05; **p<0,01 and ***p<0,001.

To evaluate the possible association of p16^High^ immune cells with a protective response against uncontrolled inflammation caused by SARS-CoV-2 in humans, we next analyzed different groups of COVID-19 patients. For this analysis, we focused on samples collected during the first two waves of pandemic back in 2020 and prior to any vaccination. Overall, the analysis was carried out in hospitalized COVID-19 patients who did not require oxygen therapy (moderate cases, n=8), those who did require oxygen therapy (severe cases, n=19) as well as in non-COVID-19 patients (n=15). All groups of patients were comparable in terms of age, sex ratio and comorbidities (Supplementary Table 2). As previously described^62,63^, patients with severe SARS-CoV-2 infection exhibited an exacerbated inflammatory response with a significant increase in IL6 (Figure 7F) and an impaired interferon response (Table 2). Analysis of PBMCs collected at the time of patient admission and before any specific treatment against SARS-CoV-2 revealed that the levels of both p16 and NNMT mRNA were significantly reduced in patients with severe COVID-19 (Figure 7I&J). These results further confirmed the potential role of p16^High^ immune cell subsets in establishing protection against severe inflammation and tissue damage induced by severe SARS-CoV-2 infection in humans.

## Discussion

The process of aging represents the gradual deterioration of organism functions with time, which affects all tissues and organs. In humans, aging is the most prevailing risk factor for the morbidity and mortality attributable to numerous acute and chronic diseases^64^. Despite relentless deterioration of our organism with age, a defined set of mechanisms has evolved to limit the negative impact of tissue damage caused by multiple factors on homeostasis. Among such protective mechanisms that rely on the concerted action of both innate and adaptive immunity as well as non-immune cells is an important but poorly understood defense strategy that limits the extent of tissue damage known as disease tolerance^24,25^. The concept of disease tolerance was originally introduced to explain additional to resistance mechanisms to fight inflammation by decreasing the host susceptibility to tissue damage without having a direct impact on the pathogens^65^. Currently, little is known about the full spectrum of tolerance mechanisms, but they seem to revolve around several evolutionary conserved stress- and damage-induced responses that confer tissue damage control in the host^65^. One of such mechanisms relies on GDF15 that among other functions acts as a strong inducer of the cell cycle arrest, senescence and is upregulated with aging^70,71^.

By exploiting different lethal conditions of inflammation and tissue damage with a focus on LPS-induced sepsis, here we show that animal survival can be significantly improved by building p16^High^ cell subsets. We argue that the mechanism of disease tolerance as the ability to withstand lethal conditions could be intimately linked with induction of a p16^High^ program in immune cells as we describe here in details but also in other cell types such as LSECs, and maintenance of a low adenosine environment (Figure 8). The latter appeared critical since reversal of a BNT2162b2 vaccine-induced downregulation of adenosine levels with dietary methionine fully blocks the development of disease tolerance in young mice while inhibiting adenosine receptors reconstitutes this protective mechanism in old animals (Figures 5L&M). In both cases, the low adenosine environment could provide optimal conditions for immune cells including Tregs and IL10-expressing macrophages to perform their immune-suppressive and tissue remodeling activities. In the case of Tregs, that we found to be one of the major p16^High^ T cell subtypes (Figure 1), it has been shown that apoptotic Tregs have much higher immunosuppressive functions than live Tregs^55^. It was found that specifically adenosine, a metabolite of extracellular ATP that is released from apoptotic Tregs, is much more immunosuppressive than typical suppressive factors such as PD-L1, CTLA-4, TGF-β, IL-35, and IL-10. In turn, reducing apoptosis through activation of a p16-dependent cell cycle arrest (Figure 5K) could limit the pool of extracellular ATP while at the same time, the NNMT-dependent consumption of SAM lowers the level of endogenous adenosine (Figure 5J and 8). This, in turn, contributes to a higher survival rate of mice in response to different types of severe inflammation and tissue damage. Indeed, we found that the presence of p16^High^ immune subsets was indispensable for animal survival in response to multiple lethal conditions such as LPS-induced sepsis (90% mice survival after induction of p16^High^ immune subsets compared to 10% in control group), acute COVID-19 infections (90% mice survival after induction of p16^High^ immune subsets compared to 30% in control group), and Ionizing irradiation (40% mice survival after induction of p16^High^ immune subsets compared to 0% in control group) (Figures 3F,G,K).

**Figure 8.**
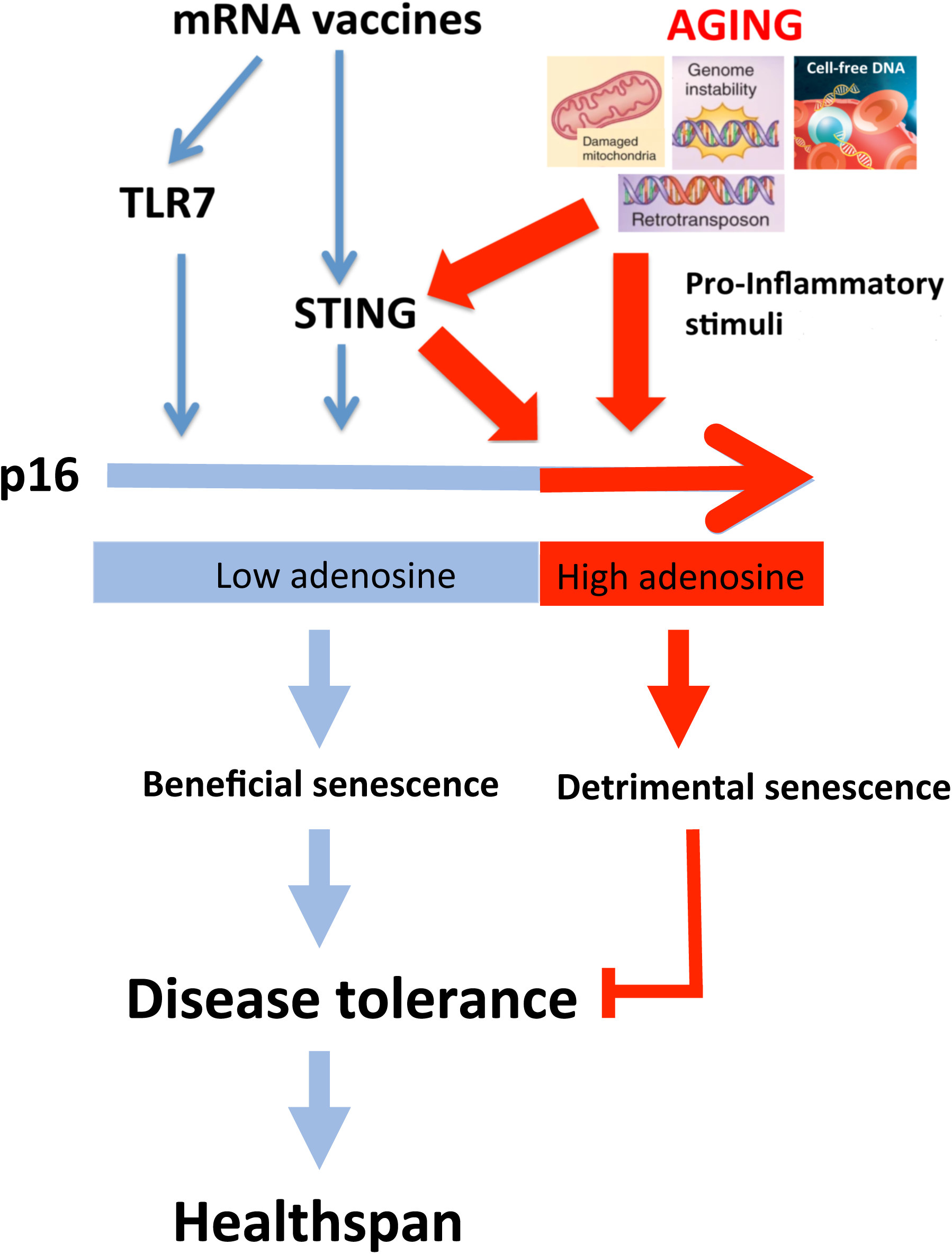
Schematic representation of disease tolerance induced by p16^High^ immune cells. Both Tlr7 and Sting pathways as long as they induce low adenosine environment (via consumption of SAM and p16-dependent cell cycle arrest and protection from apoptosis (for more details see discussion)) induce beneficial senescence that provide heighten protection from different types of inflammation and tissue damage through a mechanism called disease tolerance. In contrast, excessive inflammation contributes to detrimental senescence and derails disease tolerance.

Mechanistically, we identify TLR7 and STING as potent inducers of p16^High^ immune subsets. We further found that vaccine-based mechanism of p16^High^ immune subset induction has a strong protective effect from severe viral, bacterial and sterile inflammation. Importantly, we showed that in humans a single standard dose of vaccination with the BNT162b2 mRNA COVID-19 vaccine is sufficient to induce p16 in different immune cells. From a clinical standpoint, we found that p16-expressing immune subsets were significantly reduced in the blood of severe COVID-19 patients further supporting their critical role in preventing lethal inflammation. The clinical efficacy of the BNT162b2 mRNA vaccine in reducing the risk of hospitalization was demonstrated as early as 10 days after the first dose when the specific immune response has not yet been established^66^ in terms of development of specific antibodies^67^ and the cellular responses^68^. Our results could explain these early benefits of vaccination on the risk of hospitalization by temporarily inducing p16^high^ immune cell-dependent disease tolerance that would eventually limit the evolution towards an inflammatory form of COVID-19 thus reducing the risk of hospitalization for severe cases. Furthermore, this previously unacknowledged first line of defense in response to mRNA vaccines by building disease tolerance could play an important role in shielding against deadly pathogens while providing sufficient time to develop inhibitory antibody as lasting protection.

Our discovery of an age-induced accumulation of different p16^High^ immune cell subsets would further support the idea that it could be secondary to tissue damage and inflammation and thus, is controlled by the aged environment. Indeed, our findings that even very short-lived immune subsets as such Ly6G^+^ neutrophils contain a significant fraction of p16^High^ cells in 18-month-old mice (Figure 1A&B) indicates that such a response to the aged environment is rapid and robust. In accordance with this prediction, a recent study suggested that the aging microenvironment drives rapid epigenetic changes in macrophages based on the observation that adoptive transfer of young macrophages into old mice led to a rapid establishment of chromatin marks resembling those of aged macrophages^64^. Our findings raise the possibility that building disease tolerance by maintaining increased number of p16^High^ immune cell subsets from early age and thus before any tissue damage or inflammation could potentially provide benefits in extending healthspan, something that we now show for MDA5-deficient mice (Figure 4A&6B). It is further tempting to speculate that the mechanism of precautious activation of p16^High^ state in immune subsets in order to extend health- and lifespan is evolutionary conserved and active in long-lived animals. For example, in long-lived bats, continuous presence of multiple viruses could trigger a TLR7/STING-induced activation of a p16 program in immune cells and tissue protection against both infection and tissue damage^66^. At the same time, significantly reduced DNA-sensing in bats could keep STING activation in its low and physiological or, as also referred to, “tonic” state^67^ while preventing a strong pro-inflammatory response that could be detrimental for disease tolerance (Figures 4F&G). In another long-lived animal, the naked mole-rat, a lack of response to Poly (I:C) reported in hematopoietic stem cells may suggest a strong attenuation of MDA5-dependent sensing^68^. Importantly, as we found here that deletion of MDA5 in mice induces STING potentially as a feedback mechanism due to lack of sensing of endogenous double-stranded RNAs including from repetitive and antisense sequences^60^. This, in turn, could contribute to tonic STING activation and subsequent induction of p16 in immune subsets resulting in improvement of healthspan of the animals.

Altogether, our results provide evidence for direct and early benefits of mRNA vaccine-induced p16^High^ immune cell subsets in building disease tolerance and counteracting severe cases of viral, bacterial and sterile inflammation. In turn, sustainable activation of a p16^High^ program in defined immune subsets that could be achieved through a TLR7 and/or tonic STING activation by inhibiting MDA5 could provide a novel strategy for improving health- and potentially lifespan.

## Acknowledgements

The research in DB lab is supported by the Foundations FRM, and ANR, in BSP lab by DGOS (PHRC), ANR, Conseil Départemental des Alpes-Maritimes and Region Sud and in EH lab by the Mitsubishi Foundation and the Center for Infectious Diseases Education and Research (CiDER). The authors are very grateful to Drs. Miguel Godinho Ferreira (IRCAN, Nice) and Dr. Ana-Maria Lennon (Institute Curie, Paris) for very productive discussions. The authors acknowledge the IRCAN’s Molecular and Cellular Core Imaging (PICMI), Cytometry Core Facility (CYTOMED), Histology, Genomic and Animal Housing Facilities supported by le Cancéropole PACA, la Région PACA, le Conseil Départementale 06, l’INSERM, ARC, IBiSA, and the Conseil Départemental 06 de la Région PACA. FTM want to dedicate this work to the memory of Francisco Triana Tejas. The authors acknowledge the Lombardi Comprehensive Cancer Metabolomics Shared Resource (MSR), which is supported by Award Number P30CA051008 (P.I. Louis Weiner) from the National Cancer Institute (NIH, USA).

## Supplementary Tables

**Supplementary Table 1:** Biological characteristics of the participants in the vaccine study.

|  | Day of vaccination | D7 post-vaccination | D30 post-vaccination | P value |
| --- | --- | --- | --- | --- |
| SARS-CoV2 specific T cell response CD4<sup>+</sup> CD8<sup>+</sup> (IFN-γ, UI/ml) | 0.42 [0.27; 2.70] | 1.13 [0.91; 3.45] | 1.33 [0.50; 3.20] | 0.0375 |
| Serum IL-1β (pg/ml) | 1.32 [0.86; 5.27] | 1.86 [0.51; 8.27] | 1.06 [0.00; 2.71] | 0.2990 |
| Serum IL-6 (pg/ml) | 10.60 [4.76; 12.60] | 6.38 [1.95; 44.50] | 1.80 [1.06; 2.25] | 0.1853 |
| Serum IL-10 (pg/ml) | 1.78 [1.53; 2.33] | 1.51 [0.00; 1.77] | 4.03 [2.73; 8.07] | 0.0515 |

**Supplementary Table 2:**
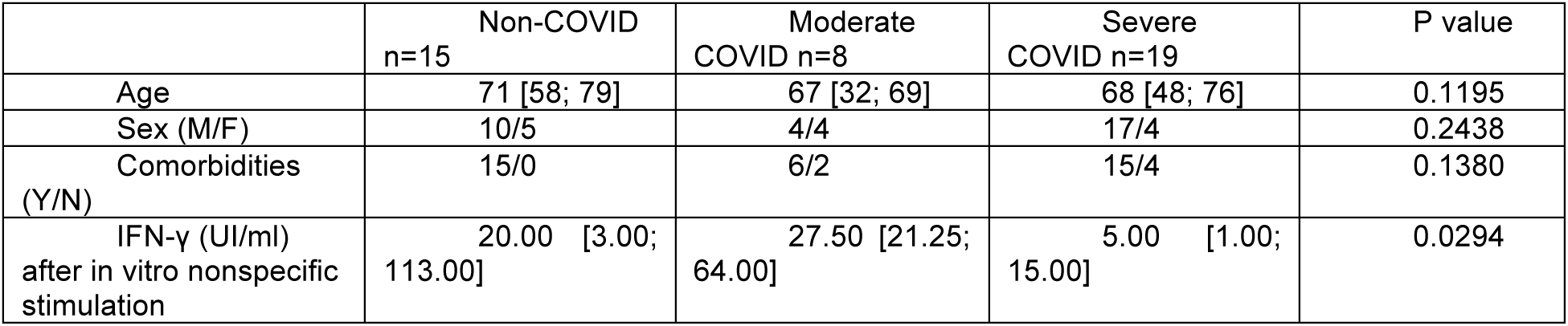
Clinical and biological characteristics of the COVID cohort.

## Material and Methods

### Animals

Previously we reported the generation of the transgenic C57BL/6 EGFP-reporter p16-Cre/R26-mTmG knock-in mice (p16-Cre/R26-mTmG henceforth). Briefly, The p16-Cre knock-in mice were generated by Ozgene (Australia) by introducing a F2A-Cre-T2A-TK-E2A-tdTomato cassette at the end of the last exon of the p16Ink4 gene. The Frt-loxed Neo cassette was removed by crossing the wt/p16-Cre heterozygous line with a homozygous FLP deleter line. Mice were then bred with Rosa26-mTmG (mixed C57BL/6J and 129/SvJ background) and Rosa26-DTA (C57BL/6 background) mice (purchased from the Jackson Laboratory). Mixed background offspring were backcrossed with C57BL/6 line to obtain p16-Cre/R26-mTmG, and p16-Cre/R26-DTA animals with pure C57BL/6 background. Experiments were performed in mixed and pure background animals. For all experiments only heterozygous mice were used for both p16-Cre knock-ins and the reporters (Rosa26-mTmG or Rosa26-DTA). For experiments, the wild type littermates, and both female and males were used. For survival experiments using lipopolysaccharides O55:B5 and γ-irradiation only C57BL/6J animals were used.

For the present study, we generated a new model of transgenic conditional knock out for Nnmt gene depending of p16 expression. A conditional knockout mouse model of Nnmt gene was generated by flanking exon 1 with loxP sites via gene targeting in mouse C57BL/6 ES cells (Ozgene, Australia). The Cre-mediated deletion of the “floxed” 1 exon after crossing with p16-Cre knock-in mice^12^ resulted in removal of the ATG coding exon specifically in p16^High^ cells. Only homozygous animals were used for experimentation. For experiments carried on this line and the wild type littermates, female and males were used.

MDA5-KO C57BL/6 mice were obtained from The Jackson Laboratory. A targeting vector was designed to delete exon1 and 400 bp of upstream sequence of the interferon induced with helicase C domain 1 *Ifih1* gene upstream sequence with a floxed MC1-neo cassette. Cre mediated excision removed the MC1-neo, leaving a single loxP site in order to abolish gene function. For experiments carried on this line and the wild type littermates, female and males were used.

For the SARS-CoV-2 (MA10) experiments wild type Balb/c females were used. Balb/c mice were purchased from SLC Japan.

### Human population

The participants of the COVID study were included from the CovImmune 1 cohort (NCT04355351) between April 2020 and September 2021 in Nice University Hospital. Participants were recruited during an emergency room consultation following COVID-19 symptoms, or as contact of a diagnosed COVID-19 case or following hospitalization for COVID-19. Patients were eligible for inclusion in this study if (i) they did not receive a vaccination regimen; (ii) SARS-CoV-2 infection was confirmed by a nasopharyngeal PCR or an antigenic test; (iii) they did not receive any COVID-19 treatment. Demographic, clinical, biological, and outcome data were collected by the study investigators and centralized in a database. Mild disease was defined as not requiring hospitalization and symptoms that did not include dyspnea and severe disease as require hospitalization and oxygen therapy. The non-COVID group is constituted of comorbid patients matched in age and sex to the COVID cohort and recruited before the start of the pandemic (March 2020) (NCT03804359).

The participants of the vaccine study were included from the CovImmune 2 cohort (NCT04429594): an epidemiological study in the context of COVID-19 that monitored periodically since July 2020 developing a SARS-CoV-2 infection or response to SARS-CoV2 vaccination. Demographic, clinical, biological data were collected by the study investigators and centralized in a database. The participants from the CovImmune 2 cohort (NCT04429594) were included to evaluate the effect of BNT162b2 mRNA COVID-19 vaccine after treatment. Seven volunteers were treated following the standard protocol approved for EMA and FDA. In total 30 μg of BNT162b2 mRNA COVID-19 vaccine were injected intramuscularly in the arm. Blood samples were taken right before vaccination and 7 and 30 days after vaccination.

These studies protocols comply with the principles of the Declaration of Helsinki and was approved by the *Comité de Protection des Personnes Sud-Ouest et Outre-Mer* institutional review board (CovImmune 1: CPP 1-20-027 ID7715 and CovImmune 2: Dossier 2-20-058 id8543). Written inform consent was obtained from all study participants.

### *In vivo* treatments

To phenocopy the effect of aging on intestinal tissue homeostasis, 2-3 months animals were treated with 2.5% dextran sulphate (M_r_ ∼40,000) (Sigma-Aldrich 42867) dissolved in drinking water for 7 days. To find the specific toll-like receptor capable to induce p16 expression, 2 months animals were injected intraperitoneally with Pam3CSK4 2 mg/kg (Invivogen tlrl-pms), Pam2CSK4 (0,1 mg/kg) (Invivogen tlrl-pm2s-1); Poly(I:C) (5 mg/kg) (Invivogen tlrl-picwlv); Lipopolysaccharides from *Escherichia coli* O55:B5 (1 mg/Kg) (Sigma-Aldrich L2880); flagellin from *Salmonella typhimurium* (0,2 mg/kg) (Invivogen tlrl-stfla); R837 (1,5 mg/kg) (Invivogen tlrl-imq); and ODN1585 (2 mg/kg) (Invivogen tlrl-1585), 48 h later, animals were euthanized for analysis. To induce severe sepsis, 2 months animals were treated with Lipopolysaccharides from *Escherichia coli* O55:B5 (40 mg/kg) (Sigma-Aldrich L2880). To induce severe tissue damage, 2 months animals were exposed to 8 Greys (Gy) of γ-irradiation. To induce lethal COVID-19 infection in mice, 10 weeks old animals were treated intranasal with the SARS-CoV-2 murine strain MA10 (4 × 10^5^ FFU), virus infections experiments were done in a biosafety level 3 facility at Osaka University. To induce an increase of p16^High^ cells in young animals, 2-3 old months animals were treated with BNT162b2 mRNA COVID-19 vaccine (5 μg by mouse) (Pfizer-BioNTech), STING activator DMXAA (10 mg/kg) (Invivogen tlrl-dmx). For TLR7 inhibition was used M5049 (1 mg/kg) (MCE, HY-134581). For STING inhibition was used H151 (10 mg/kg) (Invivogen inh-h151). For all the treatments used saline solution or PBS as a vehicle. Vehicle for each case was used as mock condition.

### Multicolor flow cytometry

#### Mouse

Mice were euthanized by cervical dislocation. Peritoneal cavity cells were collected by lavage with cold DMEM. Liver abdominal fat and lung were dissociated incubating them with 1 mg/ml collagenase type A at 37°C during 45 m and sieved consecutively with 100, 70, 40 and 30 μm strainer to obtained single cell suspensions and in the case of the abdominal fat to obtain stromal-vascular cell fraction (SVF). Spleens were mashed in cold DMEM containing DNase I at 100 μg/mL and sieved with a 75 μm and 30 μm strainer to obtain single cell suspensions. For bone marrow isolation, muscles were removed from both legs. An incision was made between lesser and greater trochanter. Bones then were centrifuged at 10,000 g during 30 seconds. Bone marrow was resuspended in HBSS containing Dnase I at 100 μg/mL and sieved with a 30 μm strainer to obtain single cells suspension. Peripheral blood was obtained mediated cheek puncture. Red blood cells were removed using lysis buffer. Cells were centrifuged at 1500 RPM during 5 minutes at 4 °C. Cells were resuspended in cytometry buffer (HBSS 1% BSA, 4% FBS, 2 mM EDTA) and stained with fluorescent conjugated antibodies (see list for specific cases) at 2 μg/mL during 25 minutes. Cells were washed 2 times with cytometry buffer. For FOXP3 staining, after membrane markers labeling, cells were fixed, permeabilized, stained and washed using the Foxp3/Transcription Factor Staining Buffer Kit (Tonbo biosciences cat. TNB-0607). Samples were analyzed in 96 wells plate by flow cytometry using Cytoflex system (Beckman-Coulter). At least 2.5X10^4^ events were recorded in singlets populations. Data were obtained and analyzed using CytExpert software. All tissues and most of the experiment were performed using the same strategy.

#### Human samples

In humans, p16 expression in T cell populations was assessed after immunostaining with specific antibody in whole blood by flow cytometry. In COVID-19 patients, flow cytometry was performed in isolated PBMC that were stored at -80C in storage solution. In whole blood samples, red blood cells were lysate with Pharm Lyse™ Lysing Buffer (BD Biosciences™). Cell surface staining was performed in 1x PBS for 30 min at 4°C. Cells were fixed and permeabilized using Transcription Factor Staining Buffer Kit (Tonbo Bioscience, San Diego, CA). Intracellular staining (FOXP3 and p16) was performed in permeabilization buffer for 30 min at 4°C. Flow cytometry data was acquired on a BD FACSLyric™ and analyzed in BD FACSuite™ software. At least 1×10^6^ events were recorded in singlets populations.

Antibodies used are listed below.

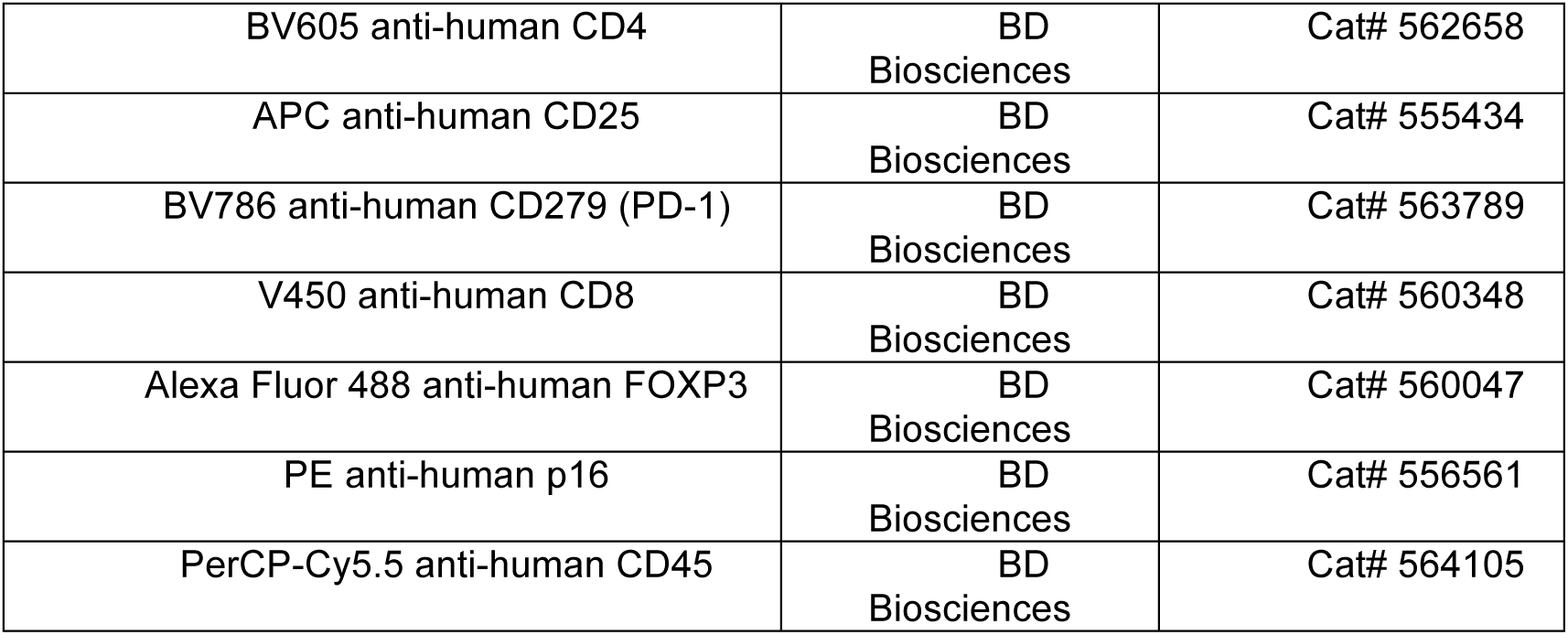
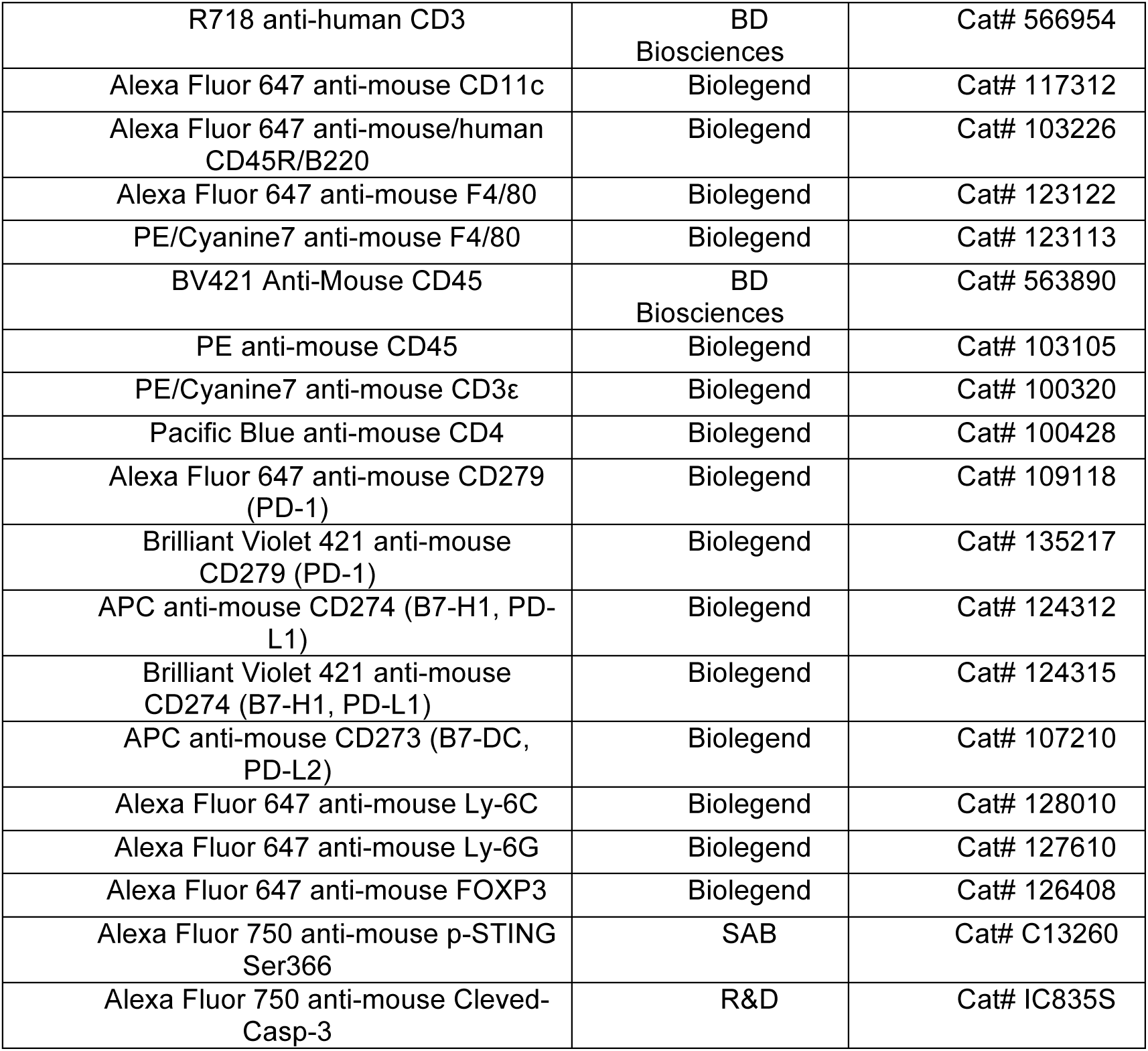

### Cell isolation and cell culture

Peritoneal macrophages were isolated using F4/80 magnetic beads (Miltenyi Biotec # 130-110-443) and LS columns (Miltenyi Biotec #130-042-401) following the manufacturés instructions. Macrophages were cultured in Roswell Park Memorial Institute medium(RPMI) 10% heat inactivated fetal bovine serum (GIBCO) and 100 U ml−1 penicillin/streptomycin (Sigma, P4333) at 37 °C with 5% CO_2_. VeroE6/TMPRSS2 cells (JCRB, 1819) were cultured in Dulbecco’s modified Eagle’s medium supplemented with 10% fetal bovine serum (MP Biomedicals, 2917354H) and 100 U ml−1 penicillin/streptomycin (Sigma, P4333) at 37 °C with 5% CO_2_. We regularly confirmed the absence of mycoplasma contamination in our cultured cells.

Healthy volunteers and COVID-19 patient PBMC were isolated by density gradient using Ficoll (Human PANCOLL, Pan Biotech™) and stored in fetal bovine serum 10% DMSO at -80°C.

### Virus preparation

The SARS-CoV-2 strain, Mouse-adapted SARS-CoV-2 (MA10), amplification and titration were previously described^69^. Briefly, titration was used as a TCID_50_ (median tissue culture infectious dose) assay by Vero/TMPRSS2 cells.

### SA-beta-Gal staining

Peritoneal cells from 2 and 12 months old p16-Cre/R26-mTmG mouse were isolated by washing peritoneal cavity with cold RPMI. Cells were incubated overnight in RPMI supplemented with 10% fetal bovine serum at 37°C and 5% CO_2_ conditions to allow them to attach to the plate. Cells were washed with PBS and fixed for 5 min at room temperature in 4% PFA, then cells were washed 3 times with PBS. Fixed cells were incubated during 8-12 hours at 37°C with staining solution containing: 40 mM Citrate-sodium phosphate pH 6; 5 mM K_3_[Fe(CN)_6_]; 5 mM K_4_[Fe(CN)_6_]; 2 mM MgCl_2_; 150 mM NaCl; and 1mg/ml X-gal.

### Fluorescence microscopy

Peritoneal cells from 12-16 months old p16-Cre/R26-mTmG mouse were isolated by washing peritoneal cavity with cold RPMI. Cells were incubated overnight in RPMI supplemented with 10% fetal bovine serum at 37°C and 5% CO_2_ conditions to allow them to attach to 10 mm slides. Subsequently, cells were fixed in 4% PFA (10 min, room temperature blocked (1 hour, 5% Goat Serum, 0.3% triton and 1%BSA) and incubated in blocking solution with first antibody (1 hour, room temperature): chicken anti-GFP (1/1800, ab13970, Abcam); rabbit anti-Lamin B1 (1/300, ab16048, Abcam); Rabbit anti-phospho-gH2AX (1/250, 9718, Cell Signalling); Rat anti-p21 (1/150, ab107099, Abcam), and Rat anti-RAD51 (1/500, BJ12112008, Bioss). The secondary antibodies used were Alexa Fluor 488 anti-chicken; Alexa Fluor 488 anti-rabbit; Alexa Fluor 647 anti-rabbit, and Alexa Fluor 647 anti-rat (1/500, Life Technology). Nuclei were counter-stained with DAPI included in mounting medium (Vectashield, H-1200). Images were taken using an HD Zeiss Axio Observer Z1 Microscope and a confocal Zeiss LSM 880 microscope (Zeiss, Gottingen, Germany) and analyzed by using ZEN (blue edition) software (Zeiss).

### Tissue microscopy

Preparation of tissue specimens for histology and immunofluorescent analysis were done as described in Grigorash et al., 2023. Briefly, tissues were fixed for the 90 minutes in ice-cold 4% PFA in PBS and passed through sucrose gradient (10-30%). After that specimens were embedded in OCT (Tissue-Tek OCT Compound, Sakura Finetek, USA) and sectioned with Cryostar NX70 cryotome (Thermo Scientific, USA). 4 mm sections were air-dried and subjected for immunofluorescent analysis following conventional protocol. Primary antibodies were used at respective concentrations: anti-CD31 (Abcam 182981; 1:50), anti-aSMA (Abcam 124964, 1:250). Secondary conjugated with AlexaFluor 647 dye anti-rabbit antibodies purchased from Thermo Scientific were used at 1:500 dilution. Samples were imaged by LSM880 confocal microscope or Axio VertA1 epifluorescent microscope (both Carl Zeiss).

### Proliferation Assay

Peritoneal cells from 12 months old p16-Cre/R26-mTmG mouse were isolated by washing peritoneal cavity with cold RPMI. Cells were incubated overnight in RPMI supplemented with 10% fetal bovine serum at 37°C 5% CO_2_ to allow them to attach to the plate. Proliferation capacity was assessed using the Click-iT® Plus EdU Assay kit (Invitrogen C10640) following the protocol provided by the manufacturer. Cells were incubated with EdU during 24 h.

### Migratory capacity Assay

Freshly isolated F4/80^+^ cells from 12 months p16-Cre/R26-mTmG mouse peritoneal cavity (Anti-F4/80 MicroBeads UltraPure, Miltenyi Order no. 130-110-443) were seeded in a 8 μm pore permeable insert (24 well plate; 3,2 mm diameter Transwells Corning) previously coated with matrigel in a migratory solution (DMEM 0,1% BSA). A lower chamber was filled with complete medium containing 100 nM Phorbol 12-myristate 13-acetate (PMA). Cells were incubated at 37°C and 5% CO_2_ during 24 h. Cells were counted using fluorescence microscope (EVOS system, Thermofisher scientific). The percentage of migrating cells were determined using cells seeded directly in 5 mm diameter wells as a control.

### Phagocytic activity Assay

Peritoneal cells from 12 months old p16-Cre/R26-mTmG mouse were isolated by washing peritoneal cavity with cold RPMI. Cells were incubated overnight in RPMI supplemented with 10% fetal bovine serum at 37°C and 5% CO_2_ to allow them to attach to the plate. Cells were incubated with beads (Invitrogen 11151D). Numbers of beads inside the cells were counted at 1, 2, 5, 10, 15 and 30 min.

### RNA sequencing

Peritoneal cavity macrophages were isolated using F4/80 beads. p16^High^ and p16^Low^ populations were separated using fluorescence activated cell sorting (FACS) (FacsAria 3 Analyzer, BD Biosciences). RNA isolation was performed using RNeasy kit (Quiagen # 74104) according to the manufacturer’s protocol. RNA degradation and contamination was monitored on 1% agarose gels. RNA purity was checked using the NanoPhotometer spectrophotometer (IMPLEN, CA, USA). RNA integrity and quantitation were assessed using the RNA Nano 6000 Assay Kit of the Bioanalyzer 2100 system (Agilent Technologies, CA, USA). A total amount of 1 mg RNA per sample was used as input material for the RNA sample preparations. Sequencing manufacturer’s recommendations and index codes were added to attribute sequences to each sample. Briefly, mRNA was purified from total RNA using poly-T oligo-attached magnetic beads. Fragmentation was carried out using divalent cations under elevated temperature in NEB Next First Strand Synthesis Reaction Buffer (5X). First Second strand cDNA synthesis was subsequently performed using DNA Polymerase I and RNase H. Remaining overhangs were converted into blunt ends via exonuclease/polymerase activities. After adenylation of 3’ ends of DNA fragments, NEBNext Adaptor with hairpin loop structure were ligated to prepare for hybridization. In order to select cDNA fragments of preferentially 150_200 bp in length, the library fragments were purified with AMPure XP system (Beckman Coulter, Beverly, USA). Then 3 ml USER Enzyme (NEB, USA) was used with size-selected, adaptor-ligated cDNA at 37 °C for 15 min followed by 5 min at 95 °C before PCR. Then PCR was performed with Phusion High-Fidelity DNA polymerase, Universal PCR primers and Index (X) Primer. At last, PCR products were purified (AMPure XP system) and library quality was assessed on the Agilent Bioanalyzer 2100 system. The clustering of the index-coded samples was performed on a cBot Cluster Generation System using SR Cluster Kit cBot-HS (Illumina) according to the manufacturer’s instructions. After cluster generation, the library preparations were sequenced on an Illumina platform and 50 bp/100 bp single-end reads were generated.

### Gene expression

Murine total RNA from peritoneal cells, liver, lung, or human PBMCs was isolated with RNeasy Mini Kit (Quiagen # 74104) according to the manufacturer’s protocol. 1 µg of total RNA was used for cDNA synthesis using RevertAid First Strand cDNA Synthesis Kit (ThermoFisher # K1621) and 30-60 ng cDNA was used for a PCR reaction. The quantitative PCR was performed with KAPA SYBR FAST qPCR Kit (KAPPA Biosystem # KR0389) using the StepOnePlus Real-Time PCR System with the following parameters: enzyme activation 2 min at 95°C, and 40 cycles of 3’’- 95°C, and 45’’- 60°C. Primers for PCR were designed using Primer-BLAST tool at NCBI with the condition of separation of primer pairs with at least one intron. The mRNA expression levels of each gene were calculated relative to β-actin expression levels. Primers sequences are listed below.

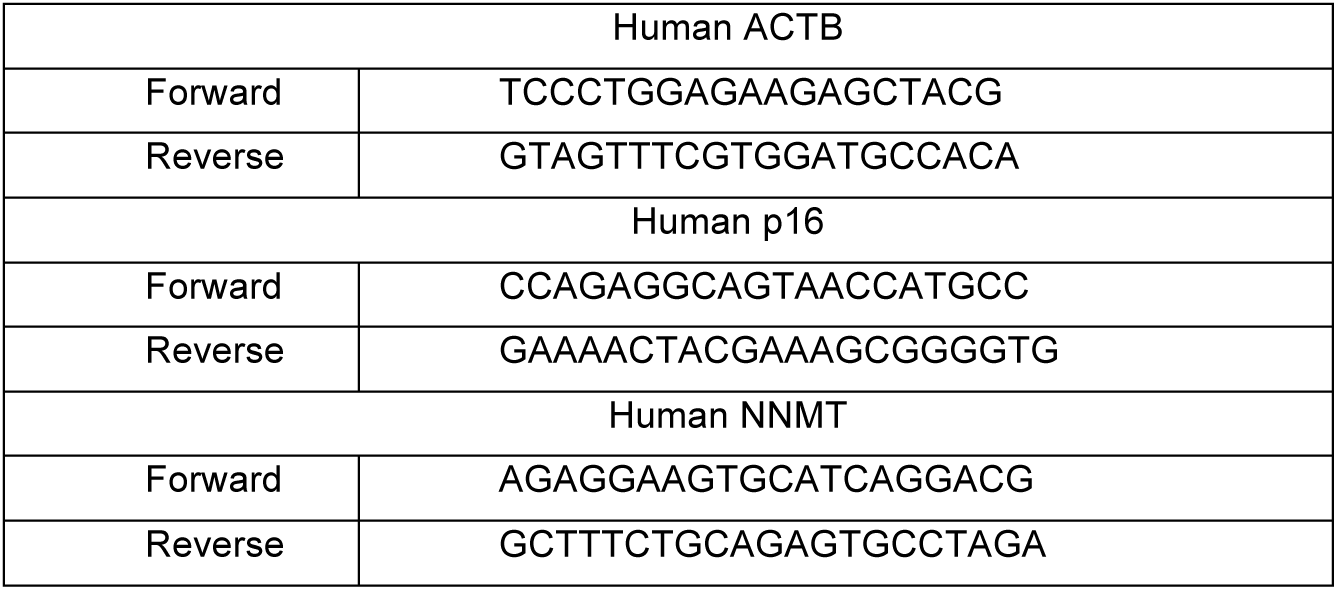
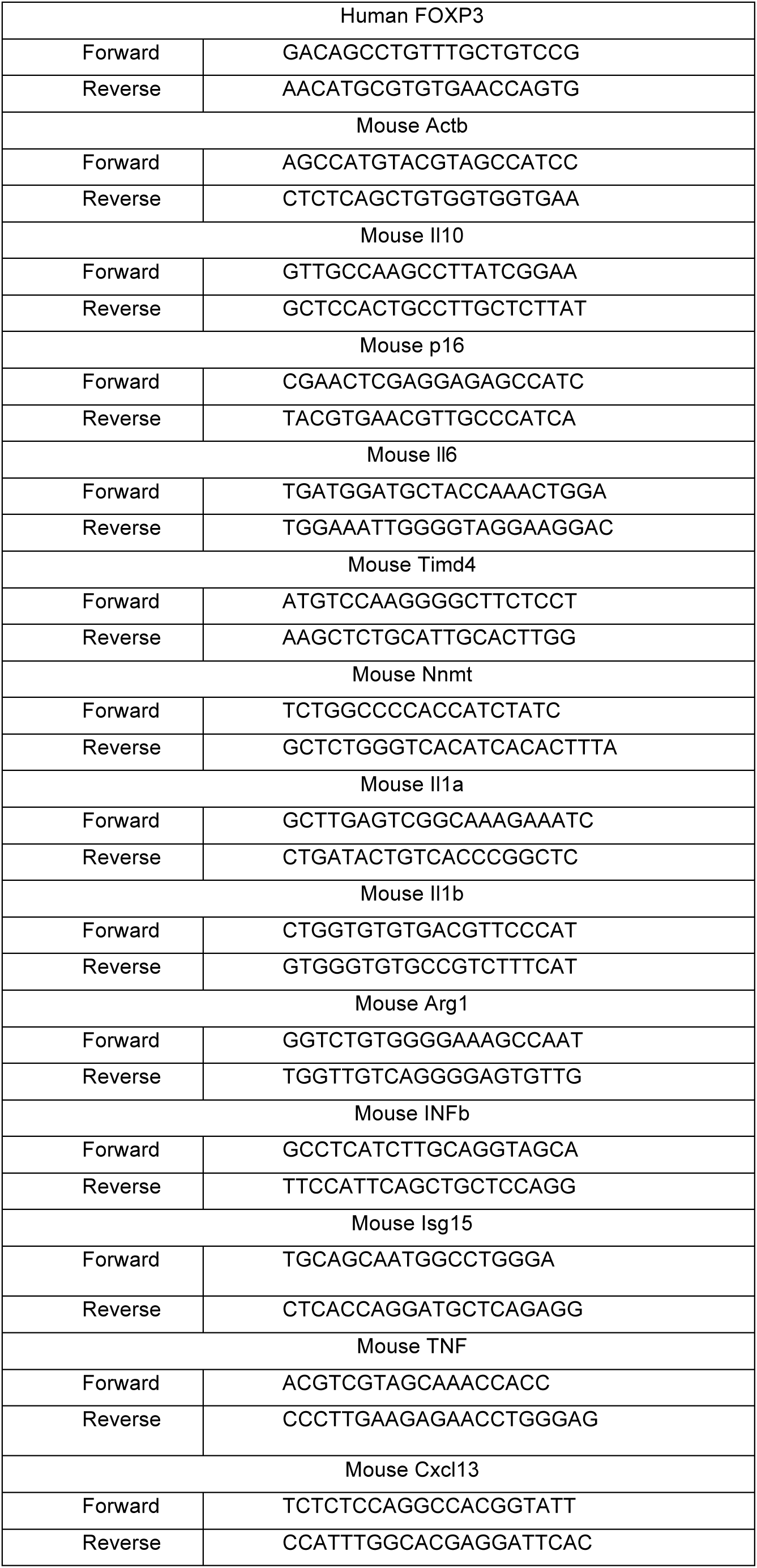

### Cytokine detection assay

Blood samples were collected for basal cytokine measure in serum. IL-6 basal level was measured in serum, with custom-designed enzyme-linked immunosorbent assay (ELISA) Multiplex Cartridges (Bio-Techne) on Automated ELISA system Ella (ProteinSimple).

### Adenosine detection assay

Enzymatic detection of adenosine was performed following the instruction provided from the supplier of the kit (Adenosine assay catalog MET-5090, CellBiolabs inc). Brefly, samples were collected in cold PBS. Peritoneal cells were lysed by sonication. Livers were disrupted with polytron plus sonication. Plasma was obtained by centrifugation (10 m 2,000 x g). Samples were assayed after being obtained or stored at -80 °C not longer than 3 months.

### Grip test analysis

Mice were placed on the food tray, which was then inverted and suspended above the home cage. The time when the animal fell was then measured (that is, latency to fall), with a threshold of 90 seconds. The test was performed in three independent sessions, three trials per session. The average performance for each session consists of the average of the three trials.

### Frailty analysis

Mice frailty were assessed using a frailty index (FI) to score relevant features observed during mice aging. The FI was obtained from the public resources in the website of Dr. Dawn Bowdish Lab, at McMaster University Hamilton, ON, Canada. (Source: http://www.bowdish.ca/lab/wp-content/uploads/2019/11/Mouse-Frailty-Scoring-2.pdf).

### RNA-Seq Data analysis

Raw data (raw reads) of fastq format were firstly processed through in-house perl scripts. In this step, clean data (clean reads) were obtained by removing reads containing adapter, reads containing ploy-N and low quality reads from raw data. At the same time, Q20, Q30 and GC content the clean data were calculated. All the downstream analyses were based on the clean data with high quality. Reference genome and gene model annotation files were downloaded from genome website directly. Index of the reference genome was built using Bowtie v2.2.3 and single-end clean reads were aligned to the reference genome using TopHat v2.0.12. We selected TopHat as the mapping tool for that TopHat can generate a database of splice junctions based on the gene model annotation file and thus a better mapping result than other non-splice mapping tools. For unigene DGE, Bowtie v0.12.9 was used to aligned single-end clean reads to the unigene sequences. HTSeq v0.6.1 was used to count the reads numbers mapped to each gene. And then FPKM of each gene was calculated based on the length of the gene and reads count mapped to this gene. FPKM, expected number of Fragments Per Kilobase of transcript sequence per Millions base pairs sequenced, considers the effect of sequencing depth and gene length for the reads count at the same time, and is currently the most commonly used method for estimating gene expression levels^58^. For unigene DGE, we use RSEM to count the reads numbers mapped to each unigene.

### Data representation and statistical analysis

Data presented based on at least three independent experiments, values are means +/- SD. For survival experiments, the sample size from all experiments is presented in each graphic. For the rest of experiments, including in vitro and in vivo, data from a representative experiment is shown. Normal distribution of data was calculated using the Shapiro-Wilk normality test. Comparison of mean values between groups was evaluated by 2-tailed Student’s t-test, Paired t-test, Wilcoxon matched pairs test, ANOVA, Tukey-Kramer test, Dunnett test, Kruskal Wallis test, Friedman test, Dunńs test, Gehan-Breslow-Wilcoxon test and Sidak’s multiple comparisons test using the GraphPad Prism program Version 8.2.1. The specific choice of test for each particular experiment is annotated in the Figure legends. p-values less than 0.05 were considered significant. Any P-value less than 0.05 was designated with one (*) asterisk; less than 0.01 with two (**) asterisks, less than 0.001 with three (***) asterisks. All p-values are reported in the Figure legends. Results are considered significant when p < 0.05. The data that support the findings of this study are available from the corresponding authors upon reasonable request. The source data underlying Figs. 1A–J, 2A–H, 3A-J, 4A-F, 5A-N, 6A-I, 7A-H will be provided as a Source Data file.

### Inclusion and ethics

These studies protocols comply with the principles of the Declaration of Helsinki and was approved by the Comité de Protection des Personnes Sud-Ouest et Outre-Mer institutional review board (CovImmune 1: CPP 1-20-027 ID7715 and CovImmune 2: Dossier 2-20-058 id8543). Data were collected by clinical researchers. Appropriate written informed consent was obtained for each participant.

All animal experiments were performed in compliance with the Animal Care and Use Committee and approved by the ethical review committee.

## References

1. Guo, J., Huang, X., Dou, L., Yan, M., Shen, T., Tang, W., and Li, J. (2022). Aging and aging-related diseases: from molecular mechanisms to interventions and treatments. Signal Transduct. Target. Ther. 7, 1–40. 10.1038/s41392-022-01251-0.

2. López-Otín, C., Blasco, M.A., Partridge, L., Serrano, M., and Kroemer, G. (2013). The Hallmarks of Aging. Cell 153, 1194–1217. 10.1016/j.cell.2013.05.039.

3. Hayflick, L., and Moorhead, P.S. (1961). The serial cultivation of human diploid cell strains. Exp. Cell Res. 25, 585–621. 10.1016/0014-4827(61)90192-6.

4. LeBrasseur, N.K., Tchkonia, T., and Kirkland, J.L. (2015). Cellular Senescence and the Biology of Aging, Disease, and Frailty. 10.1159/000382054.

5. Serrano, M., Gómez-Lahoz, E., DePinho, R.A., Beach, D., and Bar-Sagi, D. (1995). Inhibition of ras-induced proliferation and cellular transformation by p16INK4. Science 267, 249–252. 10.1126/science.7809631.

6. Serrano, M., Hannon, G.J., and Beach, D. (1993). A new regulatory motif in cell-cycle control causing specific inhibition of cyclin D/CDK4. Nature 366, 704–707. 10.1038/366704a0.

7. Hara, E., Smith, R., Parry, D., Tahara, H., Stone, S., and Peters, G. (1996). Regulation of p16CDKN2 expression and its implications for cell immortalization and senescence. Mol. Cell. Biol. 16, 859–867. 10.1128/MCB.16.3.859.

8. Baker, D.J., Wijshake, T., Tchkonia, T., LeBrasseur, N.K., Childs, B.G., van de Sluis, B., Kirkland, J.L., and van Deursen, J.M. (2011). Clearance of p16Ink4a-positive senescent cells delays ageing-associated disorders. Nature 479, 232–236. 10.1038/nature10600.

9. Di Micco, R., Krizhanovsky, V., Baker, D., and d’Adda di Fagagna, F. (2021). Cellular senescence in ageing: from mechanisms to therapeutic opportunities. Nat. Rev. Mol. Cell Biol. 22, 75–95. 10.1038/s41580-020-00314-w.

10. Tchkonia, T., Zhu, Y., Deursen, J. van, Campisi, J., and Kirkland, J.L. (2013). Cellular senescence and the senescent secretory phenotype: therapeutic opportunities. J. Clin. Invest. 123, 966–972. 10.1172/JCI64098.

11. Acosta, J.C., O’Loghlen, A., Banito, A., Guijarro, M.V., Augert, A., Raguz, S., Fumagalli, M., Da Costa, M., Brown, C., Popov, N., et al. (2008). Chemokine Signaling via the CXCR2 Receptor Reinforces Senescence. Cell 133, 1006–1018. 10.1016/j.cell.2008.03.038.

12. Grosse, L., Wagner, N., Emelyanov, A., Molina, C., Lacas-Gervais, S., Wagner, K.-D., and Bulavin, D.V. (2020). Defined p16High Senescent Cell Types Are Indispensable for Mouse Healthspan. Cell Metab. 32, 87–99.e6. 10.1016/j.cmet.2020.05.002.

13. Reyes, N.S., Krasilnikov, M., Allen, N.C., Lee, J.Y., Hyams, B., Zhou, M., Ravishankar, S., Cassandras, M., Wang, C., Khan, I., et al. (2022). Sentinel p16INK4a+ cells in the basement membrane form a reparative niche in the lung. Science 378, 192–201. 10.1126/science.abf3326.

14. Born, E., Lipskaia, L., Breau, M., Houssaini, A., Beaulieu, D., Marcos, E., Pierre, R., Do Cruzeiro, M., Lefevre, M., Derumeaux, G., et al. (2023). Eliminating Senescent Cells Can Promote Pulmonary Hypertension Development and Progression. Circulation 147, 650–666. 10.1161/CIRCULATIONAHA.122.058794.

15. Demaria, M., Ohtani, N., Youssef, S.A., Rodier, F., Toussaint, W., Mitchell, J.R., Laberge, R.-M., Vijg, J., Van Steeg, H., Dollé, M.E.T., et al. (2014). An Essential Role for Senescent Cells in Optimal Wound Healing through Secretion of PDGF-AA. Dev. Cell 31, 722–733. 10.1016/j.devcel.2014.11.012.

16. Storer, M., Mas, A., Robert-Moreno, A., Pecoraro, M., Ortells, M.C., Di Giacomo, V., Yosef, R., Pilpel, N., Krizhanovsky, V., Sharpe, J., et al. (2013). Senescence Is a Developmental Mechanism that Contributes to Embryonic Growth and Patterning. Cell 155, 1119–1130. 10.1016/j.cell.2013.10.041.

17. Muñoz-Espín, D., Cañamero, M., Maraver, A., Gómez-López, G., Contreras, J., Murillo-Cuesta, S., Rodríguez-Baeza, A., Varela-Nieto, I., Ruberte, J., Collado, M., et al. (2013). Programmed cell senescence during mammalian embryonic development. Cell 155, 1104–1118. 10.1016/j.cell.2013.10.019.

18. Omori, S., Wang, T.-W., Johmura, Y., Kanai, T., Nakano, Y., Kido, T., Susaki, E.A., Nakajima, T., Shichino, S., Ueha, S., et al. (2020). Generation of a p16 Reporter Mouse and Its Use to Characterize and Target p16high Cells In Vivo. Cell Metab. 32, 814–828.e6. 10.1016/j.cmet.2020.09.006.

19. Liu, J.-Y., Souroullas, G.P., Diekman, B.O., Krishnamurthy, J., Hall, B.M., Sorrentino, J.A., Parker, J.S., Sessions, G.A., Gudkov, A.V., and Sharpless, N.E. (2019). Cells exhibiting strong p16INK4a promoter activation in vivo display features of senescence. Proc. Natl. Acad. Sci. 116, 2603–2611. 10.1073/pnas.1818313116.

20. Hall, B.M., Balan, V., Gleiberman, A.S., Strom, E., Krasnov, P., Virtuoso, L.P., Rydkina, E., Vujcic, S., Balan, K., Gitlin, I.I., et al. (2017). p16(Ink4a) and senescence-associated β-galactosidase can be induced in macrophages as part of a reversible response to physiological stimuli. Aging 9, 1867–1884. 10.18632/aging.101268.

21. Yousefzadeh, M.J., Flores, R.R., Zhu, Y., Schmiechen, Z.C., Brooks, R.W., Trussoni, C.E., Cui, Y., Angelini, L., Lee, K.-A., McGowan, S.J., et al. (2021). An aged immune system drives senescence and ageing of solid organs. Nature 594, 100–105. 10.1038/s41586-021-03547-7.

22. Chou, J.P., and Effros, R.B. (2013). T CELL REPLICATIVE SENESCENCE IN HUMAN AGING. Curr. Pharm. Des. 19, 1680–1698.

23. Childs, B.G., Baker, D.J., Wijshake, T., Conover, C.A., Campisi, J., and van Deursen, J.M. (2016). Senescent intimal foam cells are deleterious at all stages of atherosclerosis. Science 354, 472–477. 10.1126/science.aaf6659.

24. Soares, M.P., Teixeira, L., and Moita, L.F. (2017). Disease tolerance and immunity in host protection against infection. Nat. Rev. Immunol. 17, 83–96. 10.1038/nri.2016.136.

25. Medzhitov, R., Schneider, D.S., and Soares, M.P. (2012). Disease Tolerance as a Defense Strategy. Science 335, 936–941. 10.1126/science.1214935.

26. Wang, T.-W., Johmura, Y., Suzuki, N., Omori, S., Migita, T., Yamaguchi, K., Hatakeyama, S., Yamazaki, S., Shimizu, E., Imoto, S., et al. (2022). Blocking PD-L1–PD-1 improves senescence surveillance and ageing phenotypes. Nature 611, 358–364. 10.1038/s41586-022-05388-4.

27. Sakaguchi, S., Yamaguchi, T., Nomura, T., and Ono, M. (2008). Regulatory T Cells and Immune Tolerance. Cell 133, 775–787. 10.1016/j.cell.2008.05.009.

28. Burzyn, D., Kuswanto, W., Kolodin, D., Shadrach, J.L., Cerletti, M., Jang, Y., Sefik, E., Tan, T.G., Wagers, A.J., Benoist, C., et al. (2013). A special population of regulatory T cells potentiates muscle repair. Cell 155, 1282–1295. 10.1016/j.cell.2013.10.054.

29. Arpaia, N., Green, J.A., Moltedo, B., Arvey, A., Hemmers, S., Yuan, S., Treuting, P.M., and Rudensky, A.Y. (2015). A Distinct Function of Regulatory T Cells in Tissue Protection. Cell 162, 1078–1089. 10.1016/j.cell.2015.08.021.

30. Ginhoux, F., and Guilliams, M. (2016). Tissue-Resident Macrophage Ontogeny and Homeostasis. Immunity 44, 439–449. 10.1016/j.immuni.2016.02.024.

31. Yona, S., Kim, K.-W., Wolf, Y., Mildner, A., Varol, D., Breker, M., Strauss-Ayali, D., Viukov, S., Guilliams, M., Misharin, A., et al. (2013). Fate Mapping Reveals Origins and Dynamics of Monocytes and Tissue Macrophages under Homeostasis. Immunity 38, 79–91. 10.1016/j.immuni.2012.12.001.

32. Duque-Correa, M.A., Kühl, A.A., Rodriguez, P.C., Zedler, U., Schommer-Leitner, S., Rao, M., Weiner, J., Hurwitz, R., Qualls, J.E., Kosmiadi, G.A., et al. (2014). Macrophage arginase-1 controls bacterial growth and pathology in hypoxic tuberculosis granulomas. Proc. Natl. Acad. Sci. 111, E4024–E4032. 10.1073/pnas.1408839111.

33. Campbell, L., Saville, C.R., Murray, P.J., Cruickshank, S.M., and Hardman, M.J. (2013). Local arginase 1 activity is required for cutaneous wound healing. J. Invest. Dermatol. 133, 2461–2470. 10.1038/jid.2013.164.

34. Yang, Z., Li, Q., Wang, X., Jiang, X., Zhao, D., Lin, X., He, F., and Tang, L. (2018). C-type lectin receptor LSECtin-mediated apoptotic cell clearance by macrophages directs intestinal repair in experimental colitis. Proc. Natl. Acad. Sci. U. S. A. 115, 11054– 11059. 10.1073/pnas.1804094115.

35. Man, A.L., Bertelli, E., Rentini, S., Regoli, M., Briars, G., Marini, M., Watson, A.J.M., and Nicoletti, C. (2015). Age-associated modifications of intestinal permeability and innate immunity in human small intestine. Clin. Sci. 129, 515–527. 10.1042/CS20150046.

36. Fukui, H. (2015). Gut-liver axis in liver cirrhosis: How to manage leaky gut and endotoxemia. World J. Hepatol. 7, 425–442. 10.4254/wjh.v7.i3.425.

37. Chassaing, B., Aitken, J.D., Malleshappa, M., and Vijay-Kumar, M. (2014). Dextran sulfate sodium (DSS)-induced colitis in mice. Curr. Protoc. Immunol. 104, 15.25.1-15.25.14. 10.1002/0471142735.im1525s104.

38. Fouda, A.Y., Xu, Z., Shosha, E., Lemtalsi, T., Chen, J., Toque, H.A., Tritz, R., Cui, X., Stansfield, B.K., Huo, Y., et al. (2018). Arginase 1 promotes retinal neurovascular protection from ischemia through suppression of macrophage inflammatory responses. Cell Death Dis. 9, 1–15. 10.1038/s41419-018-1051-6.

39. Edwards, J.P., Zhang, X., Frauwirth, K.A., and Mosser, D.M. (2006). Biochemical and functional characterization of three activated macrophage populations. J. Leukoc. Biol. 80, 1298–1307. 10.1189/jlb.0406249.

40. Rakoff-Nahoum, S., Paglino, J., Eslami-Varzaneh, F., Edberg, S., and Medzhitov, R. (2004). Recognition of commensal microflora by toll-like receptors is required for intestinal homeostasis. Cell 118, 229–241. 10.1016/j.cell.2004.07.002.

41. Fitzgerald, K.A., and Kagan, J.C. (2020). Toll-like Receptors and the Control of Immunity. Cell 180, 1044–1066. 10.1016/j.cell.2020.02.041.

42. Diebold, S.S., Kaisho, T., Hemmi, H., Akira, S., and Reis e Sousa, C. (2004). Innate Antiviral Responses by Means of TLR7-Mediated Recognition of Single-Stranded RNA. Science 303, 1529–1531. 10.1126/science.1093616.

43. Mantovani, S., Daga, S., Fallerini, C., Baldassarri, M., Benetti, E., Picchiotti, N., Fava, F., Gallì, A., Zibellini, S., Bruttini, M., et al. (2022). Rare variants in Toll-like receptor 7 results in functional impairment and downregulation of cytokine-mediated signaling in COVID-19 patients. Genes Immun. 23, 51–56. 10.1038/s41435-021-00157-1.

44. Rudd, K.E., Johnson, S.C., Agesa, K.M., Shackelford, K.A., Tsoi, D., Kievlan, D.R., Colombara, D.V., Ikuta, K.S., Kissoon, N., Finfer, S., et al. (2020). Global, regional, and national sepsis incidence and mortality, 1990–2017: analysis for the Global Burden of Disease Study. The Lancet 395, 200–211. 10.1016/S0140-6736(19)32989-7.

45. Li, C., Lee, A., Grigoryan, L., Arunachalam, P.S., Scott, M.K.D., Trisal, M., Wimmers, F., Sanyal, M., Weidenbacher, P.A., Feng, Y., et al. (2022). Mechanisms of innate and adaptive immunity to the Pfizer-BioNTech BNT162b2 vaccine. Nat. Immunol. 23, 543–555. 10.1038/s41590-022-01163-9.

46. Leist, S.R., Dinnon, K.H., Schäfer, A., Tse, L.V., Okuda, K., Hou, Y.J., West, A., Edwards, C.E., Sanders, W., Fritch, E.J., et al. (2020). A Mouse-Adapted SARS-CoV-2 Induces Acute Lung Injury and Mortality in Standard Laboratory Mice. Cell 183, 1070–1085.e12. 10.1016/j.cell.2020.09.050.

47. Winkler, E.S., Bailey, A.L., Kafai, N.M., Nair, S., McCune, B.T., Yu, J., Fox, J.M., Chen, R.E., Earnest, J.T., Keeler, S.P., et al. (2020). SARS-CoV-2 infection of human ACE2-transgenic mice causes severe lung inflammation and impaired function. Nat. Immunol. 21, 1327–1335. 10.1038/s41590-020-0778-2.

48. Miao, L., Li, L., Huang, Y., Delcassian, D., Chahal, J., Han, J., Shi, Y., Sadtler, K., Gao, W., Lin, J., et al. (2019). Delivery of mRNA vaccines with heterocyclic lipids increases anti-tumor efficacy by STING-mediated immune cell activation. Nat. Biotechnol. 37, 1174–1185. 10.1038/s41587-019-0247-3.

49. Berger, G., Knelson, E.H., Jimenez-Macias, J.L., Nowicki, M.O., Han, S., Panagioti, E., Lizotte, P.H., Adu-Berchie, K., Stafford, A., Dimitrakakis, N., et al. (2022). STING activation promotes robust immune response and NK cell–mediated tumor regression in glioblastoma models. Proc. Natl. Acad. Sci. 119, e2111003119. 10.1073/pnas.2111003119.

50. Decout, A., Katz, J.D., Venkatraman, S., and Ablasser, A. (2021). The cGAS– STING pathway as a therapeutic target in inflammatory diseases. Nat. Rev. Immunol. 21, 548–569. 10.1038/s41577-021-00524-z.

51. Grigorash, B.B., van Essen, D., Liang, G., Grosse, L., Emelyanov, A., Kang, Z., Korablev, A., Kanzler, B., Molina, C., Lopez, E., et al. (2023). p16High senescence restricts cellular plasticity during somatic cell reprogramming. Nat. Cell Biol. 25, 1265– 1278. 10.1038/s41556-023-01214-9.

52. Haskó, G., Linden, J., Cronstein, B., and Pacher, P. (2008). Adenosine receptors: therapeutic aspects for inflammatory and immune diseases. Nat. Rev. Drug Discov. 7, 759–770. 10.1038/nrd2638.

53. Cronstein, B.N. (1994). Adenosine, an endogenous anti-inflammatory agent. J. Appl. Physiol. 76, 5–13. 10.1152/jappl.1994.76.1.5.

54. Fredholm, B.B. (2007). Adenosine, an endogenous distress signal, modulates tissue damage and repair. Cell Death Differ. 14, 1315–1323. 10.1038/sj.cdd.4402132.

55. Maj, T., Wang, W., Crespo, J., Zhang, H., Wang, W., Wei, S., Zhao, L., Vatan, L., Shao, I., Szeliga, W., et al. (2017). Oxidative stress controls regulatory T cell apoptosis and suppressor activity and PD-L1-blockade resistance in tumor. Nat. Immunol. 18, 1332– 1341. 10.1038/ni.3868.

56. Wang, J., and Walsh, K. (1996). Resistance to Apoptosis Conferred by Cdk Inhibitors During Myocyte Differentiation. Science 273, 359–361.

57. Schmitt, C.A., Fridman, J.S., Yang, M., Lee, S., Baranov, E., Hoffman, R.M., and Lowe, S.W. (2002). A Senescence Program Controlled by p53 and p16INK4a Contributes to the Outcome of Cancer Therapy. Cell 109, 335–346. 10.1016/S0092-8674(02)00734-1.

58. Headrick, J.P. (1996). Impact of aging on adenosine levels, A1/A2 responses, arrhythmogenesis, and energy metabolism in rat heart. Am. J. Physiol.-Heart Circ. Physiol. 270, H897–H906. 10.1152/ajpheart.1996.270.3.H897.

59. Funaya, H., Kitakaze, M., Node, K., Minamino, T., Komamura, K., and Hori, M. (1997). Plasma Adenosine Levels Increase in Patients With Chronic Heart Failure. Circulation 95, 1363–1365. 10.1161/01.CIR.95.6.1363.

60. Chen, Y.G., and Hur, S. (2022). Cellular origins of dsRNA, their recognition and consequences. Nat. Rev. Mol. Cell Biol. 23, 286–301. 10.1038/s41580-021-00430-1.

61. Grunewald, M., Kumar, S., Sharife, H., Volinsky, E., Gileles-Hillel, A., Licht, T., Permyakova, A., Hinden, L., Azar, S., Friedmann, Y., et al. (2021). Counteracting age-related VEGF signaling insufficiency promotes healthy aging and extends life span. Science 373, eabc8479. 10.1126/science.abc8479.

62. Hadjadj, J., Yatim, N., Barnabei, L., Corneau, A., Boussier, J., Smith, N., Péré, H., Charbit, B., Bondet, V., Chenevier-Gobeaux, C., et al. (2020). Impaired type I interferon activity and inflammatory responses in severe COVID-19 patients. Science 369, 718–724. 10.1126/science.abc6027.

63. Reis, G., Moreira Silva, E.A.S., Medeiros Silva, D.C., Thabane, L., Campos, V.H.S., Ferreira, T.S., Santos, C.V.Q., Nogueira, A.M.R., Almeida, A.P.F.G., Savassi, L.C.M., et al. (2023). Early Treatment with Pegylated Interferon Lambda for Covid-19. N. Engl. J. Med. 388, 518–528. 10.1056/NEJMoa2209760.

64. Niccoli, T., and Partridge, L. (2012). Ageing as a risk factor for disease. Curr. Biol. CB 22, R741–752. 10.1016/j.cub.2012.07.024.

65. Martins, R., Carlos, A.R., Braza, F., Thompson, J.A., Bastos-Amador, P., Ramos, S., and Soares, M.P. (2019). Disease Tolerance as an Inherent Component of Immunity. Annu. Rev. Immunol. 37, 405–437. 10.1146/annurev-immunol-042718-041739.

66. Gorbunova, V., Seluanov, A., and Kennedy, B.K. (2020). The World Goes Bats: Living Longer and Tolerating Viruses. Cell Metab. 32, 31–43. 10.1016/j.cmet.2020.06.013.

67. Schumann, T., Ramon, S.C., Schubert, N., Mayo, M.A., Hega, M., Maser, K.I., Ada, S.-R., Sydow, L., Hajikazemi, M., Badstübner, M., et al. (2023). Deficiency for SAMHD1 activates MDA5 in a cGAS/STING-dependent manner. J. Exp. Med. 220, e20220829. 10.1084/jem.20220829.

68. Andersson, R., and Florian, M.C. (2022). Living a longer life: unique lessons from the naked mole**-**rat blood system. EMBO J. 41, e111759. 10.15252/embj.2022111759.

69. Tsuji, S., Minami, S., Hashimoto, R., Konishi, Y., Suzuki, T., Kondo, T., Sasai, M., Torii, S., Ono, C., Shichinohe, S., et al. (2022). SARS-CoV-2 infection triggers paracrine senescence and leads to a sustained senescence-associated inflammatory response. Nat. Aging 2, 115–124. 10.1038/s43587-022-00170-7.

70. Luan HH, Wang A, Hilliard BK, Carvalho F, Rosen CE, Ahasic AM, Herzog EL, Kang I, Pisani MA, Yu S, Zhang C, Ring AM, Young LH, Medzhitov R. (2019) GDF15 Is an Inflammation-Induced Central Mediator of Tissue Tolerance. Cell. 178(5):1231–1244.

71. Conte M, Giuliani C, Chiariello A, Iannuzzi V, Franceschi C, Salvioli S. (2022) GDF15, an emerging key player in human aging. Ageing Res Rev. 75. 101569.

